# Lhx9 haploinsufficiency alters transcription in the adult mouse ovary, causing subfertility and abnormal epithelium

**DOI:** 10.1101/2022.04.27.489797

**Authors:** Stephanie Workman, Megan J. Wilson

## Abstract

Understanding the molecular pathways that underpin ovarian development and function is vital for improving the research approaches to investigating fertility. Despite a significant improvement in our knowledge of molecular activity in the ovary, many questions remain unanswered in the quest to understand factors influencing fertility and ovarian pathologies such as cancer. Here we present an investigation into the expression and function of developmental transcription factor LIM Homeobox 9 (LHX9) in the adult mouse ovary. We have characterised *Lhx9* expression in several cell types of the mature ovary across follicle stages. To elucidate the function of this expression, we carried out an investigation of ovarian anatomy and transcription in a *Lhx9*^+/-^ knockout mouse model displaying subfertility. Despite a lack of gross anatomical differences between genotypes, RNA-sequencing found that 90 genes were differentially expressed between *Lhx9*^+/-^ and *Lhx9*^+/+^ mice. Gene ontology analyses revealed a downregulation of genes with major roles in ovarian steroidogenesis and an upregulation of genes with implications for ovarian cancer. Analysis of the ovarian epithelium revealed *Lhx9*^+/-^ mice have a disorganised epithelial phenotype and a significant increase in epithelial marker gene expression. These results provide an analysis of *Lhx9* in the adult mouse ovary and a new candidate for fertility research and ovarian epithelial cancer.

**Summary sentence:** *Lhx9* haploinsufficient mice are subfertile with altered expression of steroid genes in the adult ovary and abnormal ovarian surface epithelium.

## Introduction

While all organs in the adult mammalian system require a certain level of plasticity, cellular renewal, tissue growth and change, the ovary is more complex and dynamic than many. Through the development and maintenance of follicles, the cell populations in the ovary are constantly changing during the oestrus cycle. This requires a genetic landscape that can quickly switch to promote cellular growth and proliferation when needed, achieved through precise intercellular signalling and response to endocrine activity^1–3^. The ovary must also undergo rapid remodelling and repair following the rupture of the ovarian surface epithelium during ovulation^4^. Unfortunately, the current understanding of how these genetic systems work in the ovary is limited. Despite a good grasp of the physiological and anatomical changes across the oestrous cycle, the molecular regulators of these functions remain uncertain.

A handful of marker genes distinguish oocytes from somatic cells and between several somatic populations^1^. Additionally, several genes have been identified as new markers of cell populations resulting from single-cell sequencing studies^5–7^. Despite these new technologies, the intricacies of the ovary make it challenging to assign genes to expression and function. For example, shifting cell populations during follicle maturation. In vivo analysis of ovarian dynamics can be more complex than other organ systems. Currently, there is no in vitro organoid model that can replicate all of the complexity of ovarian biology, although this seems likely to soon change^8,9^. As a result, the underlying causes of most infertility, subfertility, and other reproductive pathologies such as ovarian cancers remain a mystery. To understand these pathologies on a genetic level, the selection of genes we investigate should be expanded to study better those whose roles and functions are poorly understood. This is necessary clinically^10^ and also in increasing the scope of reproductive research.

An excellent place to start is with genes that play a role in developing the earliest ovarian precursor tissue in mammals, the genital ridge. LIM Homeobox 9 (*Lhx9*) is a transcription factor with wide roles during vertebrate development. Part of the LIM family of genes influences the development of the brain, heart, limbs, and gonad by regulating the expression of several genes^11–13^. Despite this, a complete knockout of *Lhx9* in the mouse results in no severe phenotypes in any tissue except the gonad, which displays complete agenesis due to reduced somatic cell proliferation in the genital ridge ^14^. Furthermore, Birk *et al*. showed a reduced expression of Nuclear Receptor subfamily 5 group A1 (*Nr5a1*) in the *Lhx9*^-/-^ genital ridge at E11.5, indicating that loss of *Lhx9* function led to altered transcription of essential developmental genes. Previously, *Lhx9* has shown a peak expression in the gonad at E13.5, decreasing until birth^15^. In addition, there has been some reporting of Lhx9 expression in the postnatal ovary, although this has not been investigated thoroughly and is seen only upon an investigation of supplementary materials^7,16,17^. Thus, we aimed to characterise the expression of *Lhx9* in the adult mouse ovary and determine its potential function as a fertility factor.

Here, we describe the expression of *Lhx9* mRNA and protein in multiple cell types of the adult mouse ovary. Using an *Lhx9* knockout mouse line, we have shown a premature reduction in fertility of *Lhx9* heterozygotes compared to wildtype littermates. RNA sequencing of the postnatal *Lhx9*^+/-^ ovary has revealed differential expression of several genes, including those involved in ovarian steroidogenesis, epithelial regulation, and carcinogenesis. Together, these results expand the known role of *Lhx9* beyond the scope of ovarian development and raise questions about the role of developmental transcription factors in adult tissues.

## Results

### *Lhx9* is expressed in the adult mouse ovary

To determine the role of *Lhx9* in the adult ovary, we aimed to first characterise the spatial expression of *Lhx9* via in situ hybridisation (ISH) and immunohistochemistry (IHC). Ovaries from mice aged five months and in the proestrus stage of the oestrous cycle were used. ISH was carried out using a probe capturing all five *Lhx9* isoforms^12^. *Lhx9* mRNA expression was seen in several cell types across all follicle stages from primary to late antral (Fig. 1A). Expression of *Lhx9* mRNA was strong in the oocytes and surrounding granulosa cells of all follicle stages, including the cumulus and mural granulosa cells of the larger follicles (Fig. 1A). There was a small amount of staining in some luteal cells throughout the ovary, although this was not as strong as in the other stained somatic populations (Fig. 1A). *Lhx9* mRNA expression was also seen in parts of the ovarian surface epithelium (Fig. 1A). There was seemingly no pattern to the location of epithelial expression, but it was seen across the entire ovary and multiple sections.

**Figure 1:**
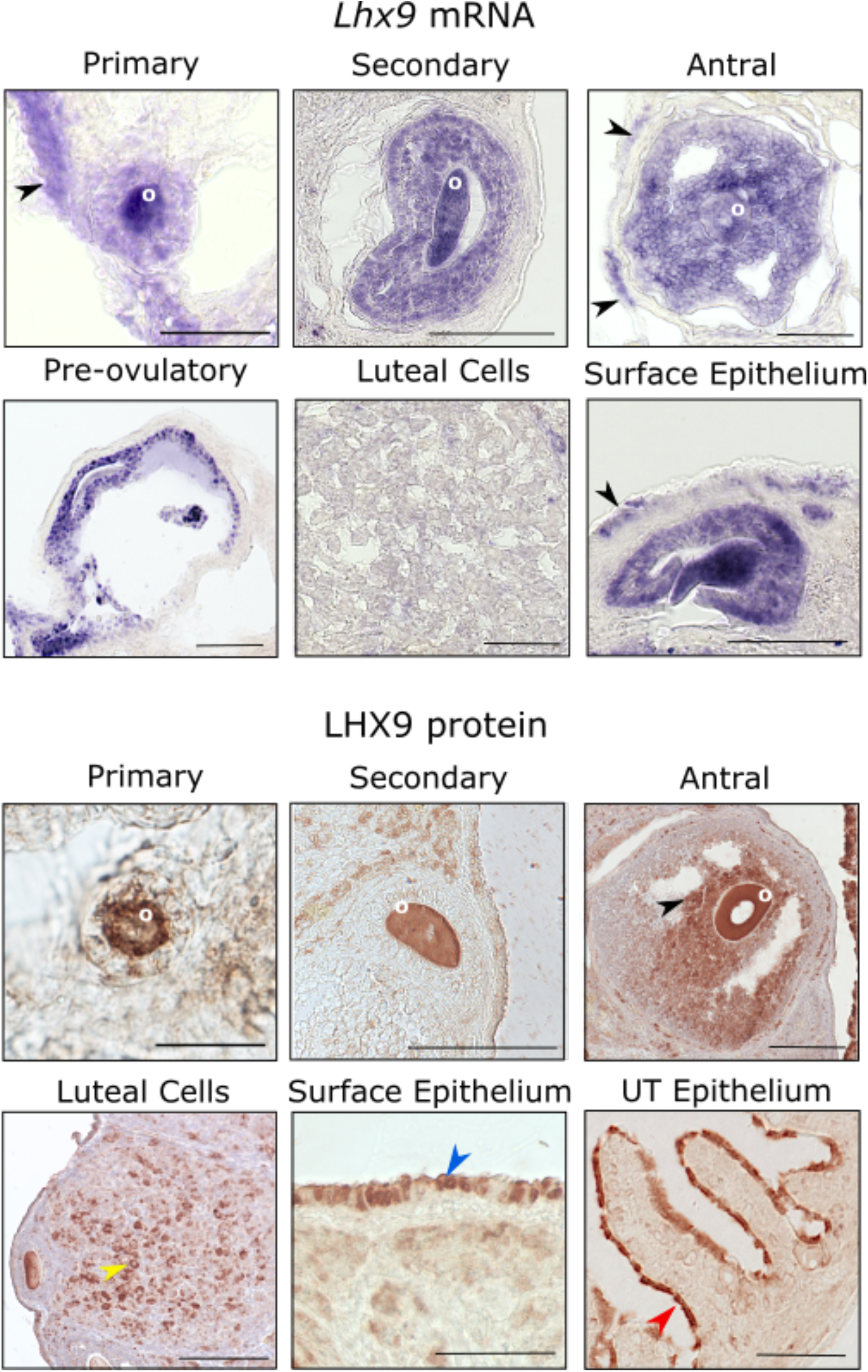
*Lhx9* mRNA and protein expression in the 5-month-old proestrus wildtype mouse ovary detected by *in situ* hybridisation and immunohistochemistry. **A.** *Lhx9* mRNA was detected across all follicle stages and selected somatic populations. Primary and secondary follicles show strong staining in the oocytes and some staining in the surrounding granulosa cells. Staining was seen in the oocytes (antral and large pre-ovulatory follicles) and the cumulus and mural granulosa cells. Luteal cells show some relatively minor staining compared to the somatic granulosa cells. Some regions of the ovarian surface epithelium were stained (black arrowheads) but not consistently across the entire surface epithelium. **B.** LHX9 protein was detected in the oocytes of primary and secondary follicles. The antral follicle staining was also seen in the oocyte and seen strongly in the cumulus granulosa cells (black arrowhead). Some cells of the corpora lutea show staining (yellow arrowhead). Similarly, the mRNA select regions of the ovarian surface epithelium were stained (blue arrowhead). The uterine tube epithelium was stained positive for LHX9 in most cells (red arrowhead). Sale bar = 100 μm except for primary follicle images, where the scale bar = 20 μm. o = oocyte.

The expression of LHX9 protein in the ovary was detected via IHC. Like the mRNA, protein expression was seen across several follicle stages from primary to late antral (Fig. 1B). The staining in the oocytes was still intense; however, the protein expression in the early-stage follicle granulosa cells appeared to be lesser than the mRNA. In later stage follicles, granulosa cell expression was more comparable with the mRNA. Protein expression was localised mainly to the cumulus granulosa cells in antral follicles (Fig. 1B). There was additionally clear expression in some of the theca cells surrounding the larger follicles. As with the mRNA, LHX9 protein was staining detected in some cells of the corpora lutea and regions of the ovarian surface epithelium (Fig. 1B). There was also expression in the luminal uterine tube epithelium; however, expression was in most epithelium cells in contrast to the ovarian surface epithelium, where expression was only in a few select regions (Fig. 1B).

### *Lhx9* haploinsufficiency does not impact folliculogenesis

Considering the specific expression pattern of *Lhx9* observed in the adult ovary, we aimed to examine the impact of reduced *Lhx9* on the ovary anatomically. The reduced proliferation of somatic cells observed by Birk et al. in *Lhx9*^-/-^ mice resulted in complete gonadal agenesis^14^. Therefore, we used *Lhx9* heterozygotes (*Lhx9*^+/-^) for all analyses (Fig. 2A). We first confirmed that the protein level of LHX9 was significantly reduced in the *Lhx9*^+/-^ gonads by Western blot (Fig. 2B; p < 0.0001). Recording the number of pups per litter over time revealed that *Lhx9*^+/-^ mice had significantly reduced pups per litter after five months, decreasing rapidly at around four months (Fig. 2C; p = 0.0042). At about seven months, these mice became infertile much sooner than their wildtype counterparts.

**Figure 2:**
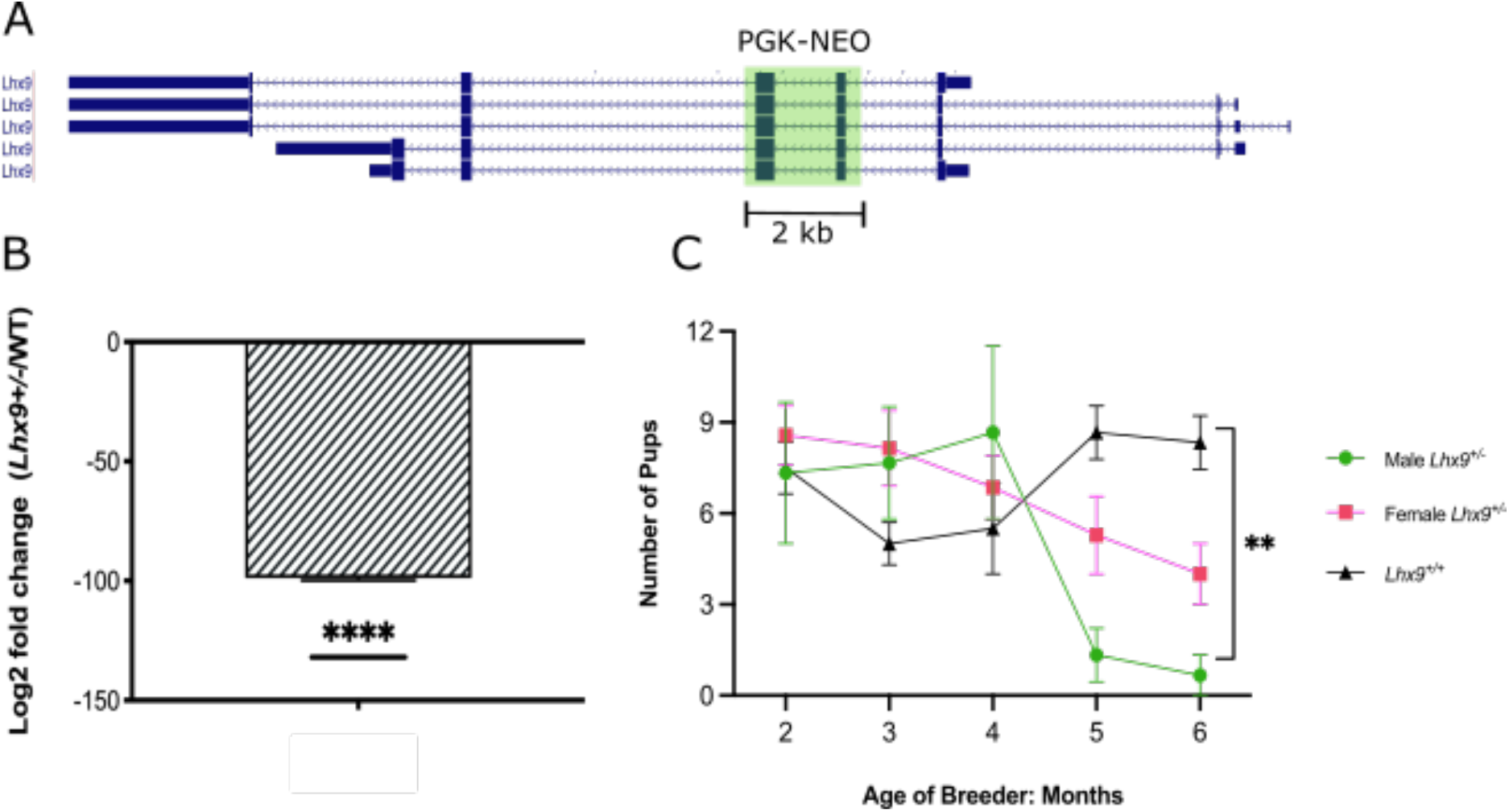
Generation and analysis of *Lhx9^+/-^* mice. A: UCSC Genome Browser view of the *Lhx9* isoforms. Exons two and three highlighted in green show the insertion site of the PGK Neomycin cassette interrupting the protein. B: Lhx9 protein is significantly decreased in the *Lhx9*^+/-^ mice as measured by Western blot. C: Litter size over time is significantly reduced in *Lhx9*^+/-^ mate pairs.

To investigate this observed subfertility, we carried out a histological analysis of the *Lhx9*^+/-^ ovary structure, including counting the large follicles and corpora lutea. There were no immediately apparent differences in the general structure or follicle morphology between the *Lhx9*^+/+^ and *Lhx9*^+/-^ ovaries (Fig. 3). The heterozygote ovary contained corpora lutea and follicles across the range of stages of development (Fig. 3D-F). A count of the antral, pre-antral follicles and corpora lutea revealed no difference in the number of significant structures between the genotypes (Fig. 3G-I; p = 0.18, 0.65 and 0.49 respectively).

**Figure 3:**
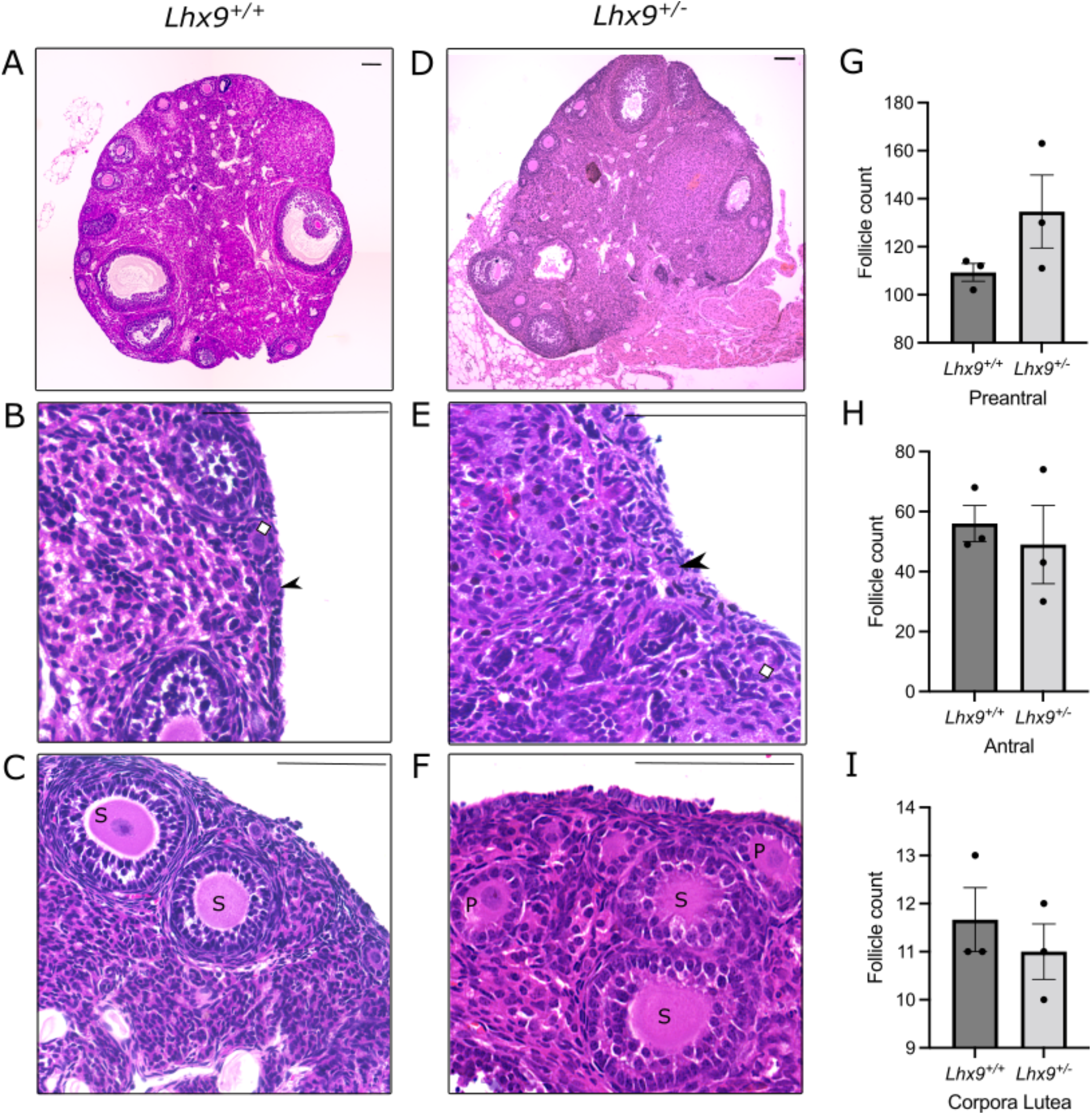
*Lhx9*^+/-^ ovaries have no obvious structural differences in large follicle numbers than *Lhx9*^+/+^ ovaries. (A, B, C). Wildtype *Lhx9*^+/+^ ovaries stained with H&E show ovarian structures (D, E, F). Heterozygous *Lhx9*^+/-^ovaries stained with H&E show normal ovarian structures like that of the wildtype control. (G, H, I) Preantral, antral and corpora lutea show no significant differences in number between *Lhx9*^+/+^ and *Lhx9*^+/-^ ovaries. Scale bars = 100 μm.

### RNA Sequencing of *Lhx9*^+/-^ ovaries

To gain insight into the transcriptional landscape of the *Lhx9*^+/-^ ovary, we carried out RNA sequencing of the five-month proestrus *Lhx9*^+/+^ and *Lhx9*^+/-^ ovaries, just prior to fertility reduction, followed by differential gene expression analysis. Approximately 26 M reads were generated for the six samples. Between 77.2% and 89.7% of genes were mapped for 19518262 – 22967318 uniquely mapped reads. A total of 66 genes were upregulated, and 24 genes downregulated in the *Lhx9*^+/-^ ovary following combined analysis with EdgeR and DESeq2 (Fig. 4A-E, File S1). The top ten significant genes with the highest fold change in either direction (following combined analysis) are shown in Figure 4D. A selection of DEGs was used to validate the RNA-sequencing results through RT-qPCR (Fig. 4F and G). A Pearson’s r-value of 0.8525 and r^2^ score of 0.7268 confirmed the differentially expressed genes validated in independent samples (n = 3/genotype).

**Figure 4:**
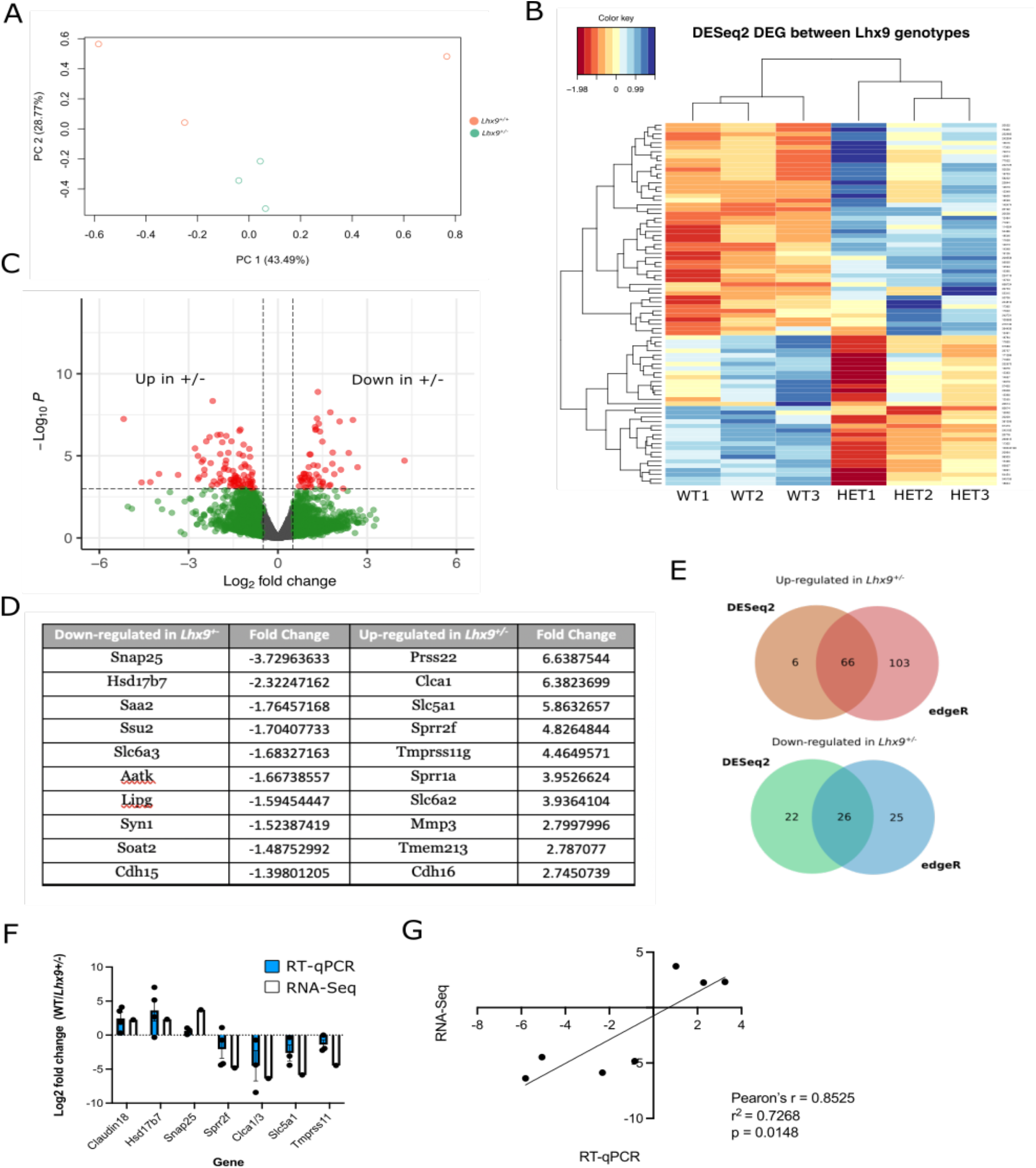
Identification and validation of differentially expressed genes between *Lhx9*^+/+^ and *Lhx9*^+/-^ ovaries. (A) Principal component analysis *Lhx9*^+/+^ samples (orange) and *Lhx9*^+/-^ samples (green) separated by principal component analysis. (B) Heat map of the differentially expressed genes generated by EdgeR with individual hierarchical clustering of *Lhx9*^+/+^ and *Lhx9*^+/-^ biological replicates. (C) Volcano Plot of significantly DEGs between *Lhx9*^+/+^ and *Lhx9*^+/-^ ovaries generated by analysis with DESeq2 Red dots represent significantly differentially expressed genes with a log fold change of < −1 or > 1. Points to the left indicate genes upregulated in the *Lhx9*^+/-^ ovary. Points to the right indicate genes downregulated in the *Lhx9*^+/-^ ovary. (D) Table of the top 10 DEGs both upregulated in *Lhx9*^+/-^ and downregulated in the *Lhx9*^+/-^. Fold changes are from the EdgeR data. (E) Venn diagram showing the shared differentially expressed genes between *Lhx9*^+/+^ and *Lhx9*^+/-^ ovaries generated by analysis with EdgeR and DESeq2. 66 upregulated and 26 downregulated genes were shared in both analyses. (F) Bar plot showing the Log2 fold change of RNA Sequencing results and the associated RT-qPCR results (n=3). (G) Correlation plot of the RNA-seq and RT-qPCR results and associated Pearson’s r and r^2^ values show a significant correlation between the two datasets. The line represents a linear regression.

The normalised read counts of crucial marker genes were compared between the genotypes to find changes in the cellular populations of the ovary as a result of reduced *Lhx9* (Fig. 5). As expected, given the results of follicle counting, no difference in mRNA expression was observed for the germ cell marker *Ddx4/Vasa* (padj = 0.66). The major granulosa cell markers *Foxl2* and *Fst* showed no significant differences between genotypes (Fig. 5, padj = 0.06 and 0.69, respectively). However, this doesn’t account for different granulosa populations associated with follicles at various stages of growth. The ovarian surface epithelial marker *Prom1* had 1.3-fold increased expression in the *Lhx9*^+/-^ compared to the wildtype ovary (Fig. 5; padj = 0.004). The transcript expression of two stromal and theca cell markers, *Lum* and *Cyp11a1*, were significantly altered in the heterozygote (Fig. 5, padj = 0.047 and 0.032, respectively). *Lum* expression increased 0.9-fold in the *Lhx9*^+/-^ovary, and *Cyp11a1* mRNA expression reduced 1.1-fold (File S1).

**Figure 5:**
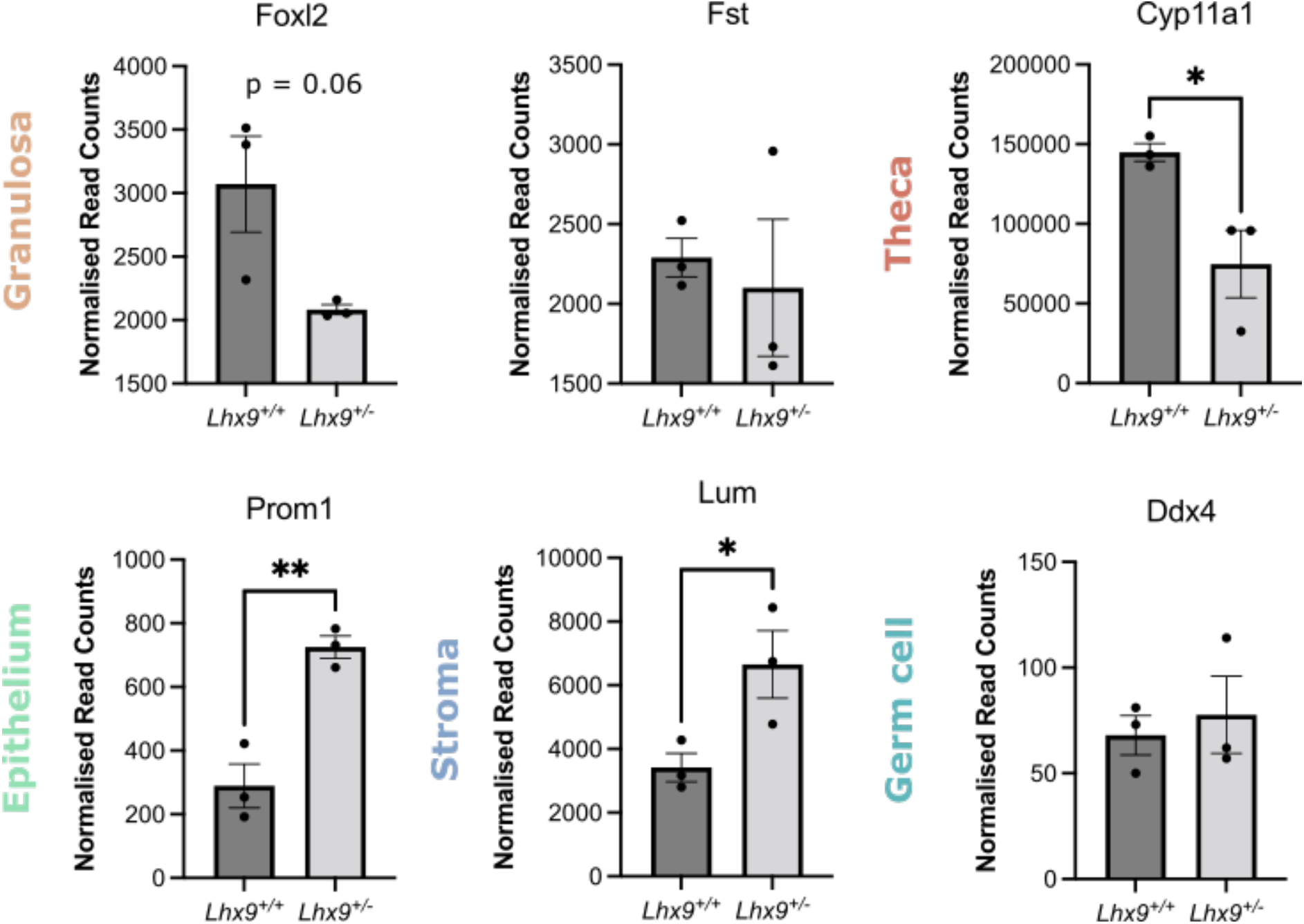
Normalised read counts of marker genes for ovarian somatic cell populations in *Lhx9*^+/+^ and *Lhx9*^+/-^ovaries. Granulosa cell markers *Foxl2* and *Fst* had no significant difference in read counts but showed a trend toward decreased expression in *Lhx9*^+/-^. The theca cell marker *Cyp11a1* had significantly fewer read counts in the *Lhx9*^+/-^ ovaries. Ovarian surface epithelium marker *Prom1* and stromal marker *Lum* had significantly more read counts in the *Lhx9*^+/-^ ovary than the *Lhx9*^+/+^. Germ cell marker *Ddx4* showed no significant difference. Statistical significance calculated by unpaired t-test. * p <0.05, ** p < 0.01.

Gene Ontology analysis of the genes with reduced mRNA expression in the heterozygote revealed their involvement in several pathways and processes linked to ovarian function. Analysis of overrepresented biological processes including ovarian steroids and cholesterol (Fig. 6A; File S2). KEGG and Panther database analysis also identified several pathways relating to ovarian steroids, including many relating to the metabolism and synthesis of cholesterol, an essential precursor for steroidogenesis (Fig. 6B). Primarily these terms are representative of decreased *Lss, Lipg, Hsd17b7*, and *Tm7sf2* gene expression, all genes involved in cholesterol and steroid metabolism pathways^18–20^. Finally, analysis of over-represented mammalian phenotypes using the MGI database again highlights lipid and steroid-related terms (Fig. 6C).

**Figure 6:**
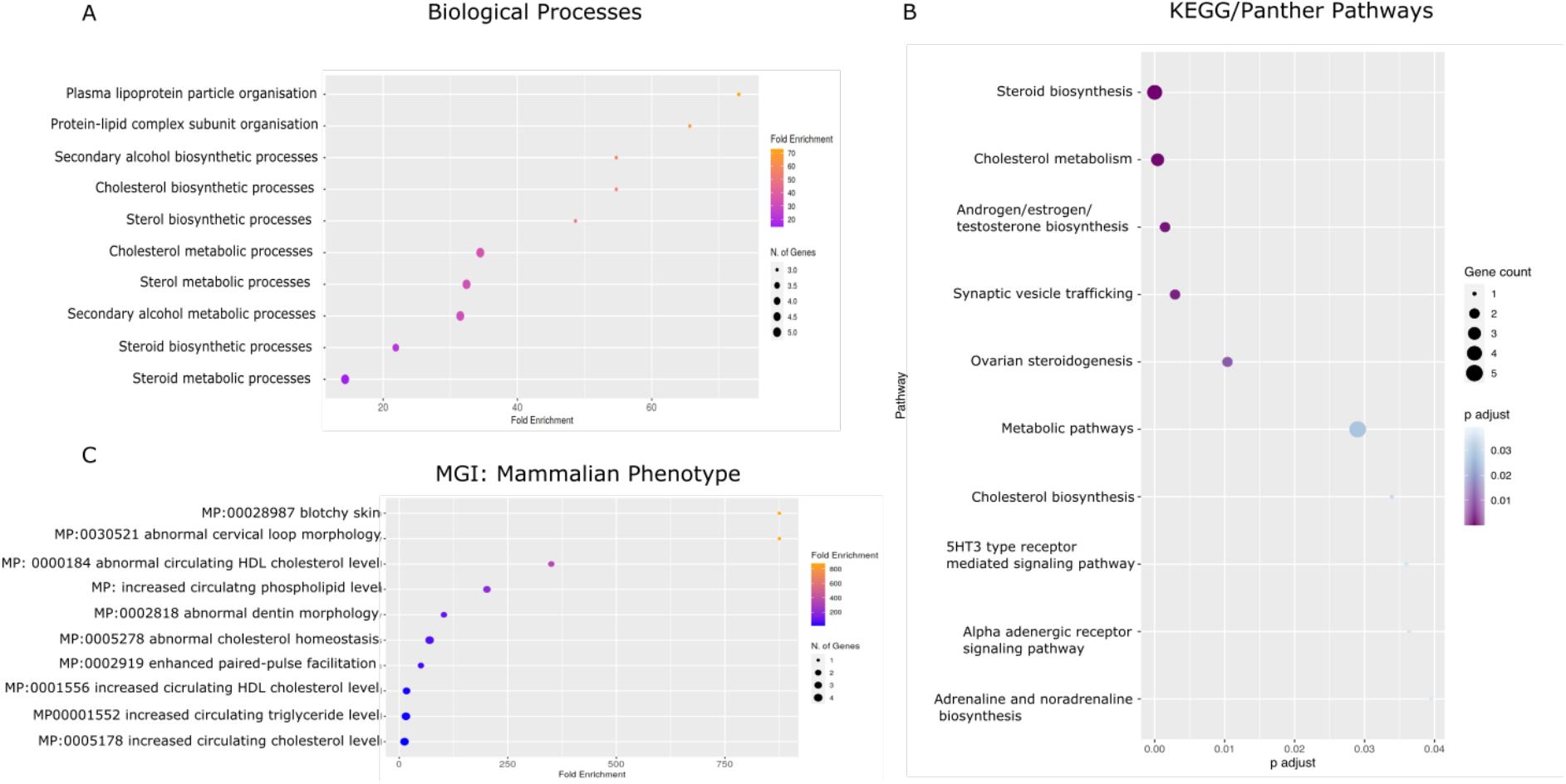
Gene Ontology analysis of genes downregulated in the *Lhx9*^+/-^ ovary. A. Over-represented biological processes in the gene list identified using ShinyGO. B. Pathways are over-represented in the gene list identified from the KEGG and Panther databases using the KOBAS software. C. Mammalian phenotypes are over-represented in our gene list identified in the MGI database using ShinyGO.

A similar analysis of the genes upregulated in the heterozygote showed several biological processes associated with epithelium differentiation and development. (Fig. 7A; File S2). Enriched pathways in the KEGG and Panther databases include transcriptional misregulation in cancer and several terms related to cytochrome P450 (Fig. 7B). In addition, analysis of tissues and cell types over-represented in the gene list identified two related epithelial tissues, glandular and uterine, and cancer stem cells (Fig. 7C).

**Figure 7:**
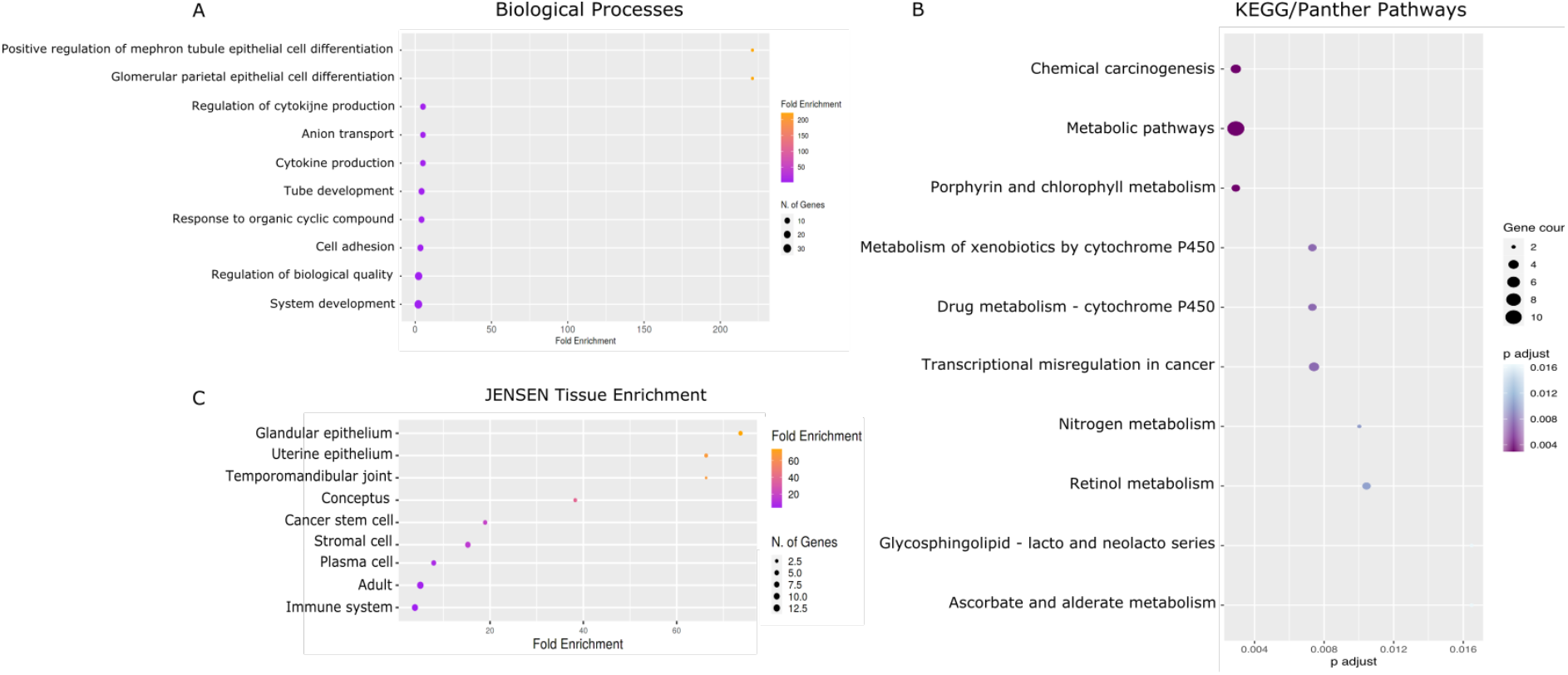
Gene Ontology analysis of genes upregulated in the *Lhx9*^+/-^ ovary. A. Over-represented biological processes in the gene list identified using ShinyGO. B. Pathways are over-represented in the gene list identified from the KEGG and Panther databases using the KOBAS software. C. Tissues and cell types enriched in the gene list identified in the JENSEN tissue enrichment database using ShinyGO.

### *Lhx9* impact on the ovarian surface epithelium

Following the identification of several epithelia related GO terms in the genes upregulated in the heterozygote (Fig. 7), together with the altered expression of epithelial marker genes (Fig. 5), we decided to further investigate the ovarian surface epithelium (OSE) genes. The OSE is an incredibly dynamic part of the ovary that must undergo rapid proliferation and repair following ovulation. The presence of a small group of stem-like cells that turn over to maintain the epithelial population has been confirmed through lineage tracing experiments^21^. This level of proliferative capacity may be why the OSE is implicated in the development of 90% of ovarian carcinomas^22^.

Analysis of the upregulated DEG list with the ChIP Enrichment Analysis (ChEA) database^23^ found enrichment for known gene targets of multiple transcription factors linked to epithelial proliferation in ovarian carcinomas, including SUZ12, EZH2, TP53, and JARID2 (Fig. 8), some of these proteins form the Polycomb Repressor Complex ^24–27^.

**Figure 8:**
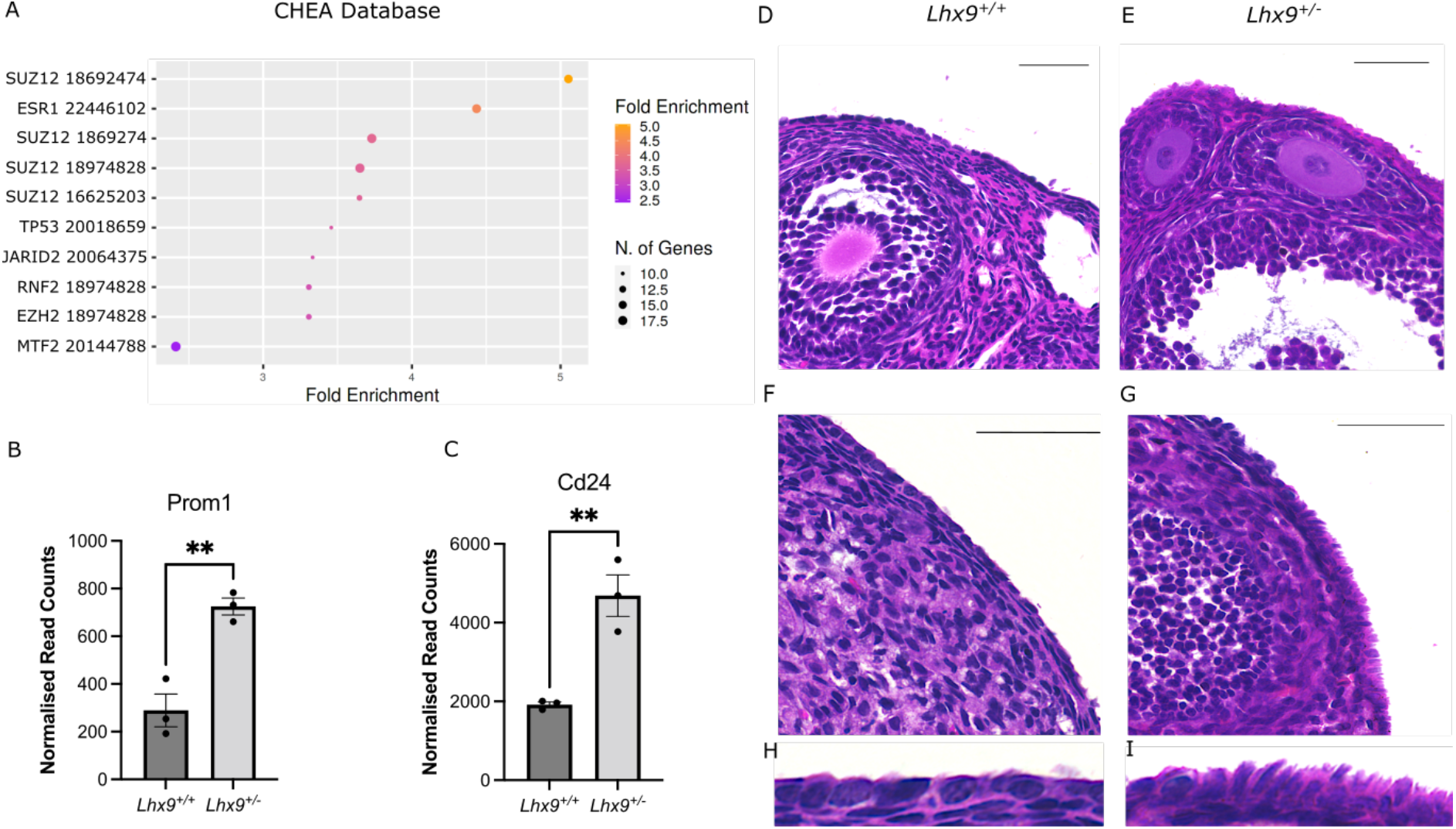
*Lhx9*^+/-^ mice have altered expression of epithelial markers and abnormal ovarian surface epithelium. (A) Analysis of transcription factors with targets overrepresented in the upregulated gene list. Numbers refer to specific gene set identities in the database from individual experiments. Identified using the Chea database via ShinyGO. (B) Normalised read counts of ovarian surface epithelial marker *Prom1* show a significant increase in expression in the *Lhx9*^+/-^ ovaries. Ovarian carcinoma marker *Cd24* also shows a significant increase in the heterozygotes (D, F, H). Wildtype *Lhx9*^+/+^ ovaries display normal, flattened cuboidal epithelium. (E, G, I). *Lhx9*^+/-^ ovaries display a highly variable epithelium, including densely packed pseudostratified cuboidal and columnar cells. Scale bars = 100 mm. ** = p <0.01.

In addition to a significantly higher read count for epithelial marker *Prom1*, the ovarian carcinoma marker *Cd24* also showed substantially greater read counts in the *Lhx9*^+/-^ ovary^28^ (Fig. 8 C, Padj = 0.0065). Both genes appeared in the significantly enriched GO terms “positive regulation of nephron tubule epithelial cell differentiation” and “glomerular parietal epithelial cell differentiation” (Fig. 7; File S2). Additionally, four genes identified as being part of the KEGG pathway “Transcriptional misregulation in cancer” (File S2) were upregulated in the heterozygous ovary, three of which have been shown to have higher expression in ovarian carcinomas, *Mmp3, Hoxa10,* and *Prom1* ^6–28^.

Analysis of the epithelium in the H&E-stained ovaries showed some striking differences between the genotypes (Fig. 8 D-I). While the epithelium in a normal ovary does show some variation, it comprises flattened cuboidal cells, with a few small regions of columnar epithelium, usually following follicle rupture. This was reflected in the *Lhx9*^+/+^ samples (Fig. 8 D, F and H). The *Lhx9*^+/-^ ovaries showed a significant level of variation, with some regions displaying a similar flattened cuboidal shape but others comprising cells with a more columnar or pseudostratified cuboidal appearance. These cells often appeared less organised and more densely packed, giving an overgrown appearance (Fig. 8 E, G and I).

## Discussion

### Examining Lhx9 expression to inform function

*Lhx9* mRNA and protein expression in the adult ovary was detected in the somatic cells and oocytes of follicle stages from primary to pre-ovulatory. Analysis of *Lhx9* expression in the microwell sequencing expression data at the Mouse Cell Atlas^32^ supports this expression pattern, finding *Lhx9* in epithelial, cumulus, and luteal cells (Fig. S1). There was no cluster for oocyte expression, likely due to the difficulties of capturing oocytes in single-cell data due to size exclusion factors. Thus, we were unable to confirm this expression using this database. Despite this, Lhx9’s consistent expression in the oocyte is intriguing, considering this expression is not seen during embryonic development or in the neonatal ovary^14,33^. The closely related gene, *Lhx8*,a known critical fertility factor^34^, is expressed in the oocytes of developing and adult ovaries. While there is a relatively high level of redundancy within the LIM Homeobox gene family, this has not been shown for *Lhx9* and *Lhx8* in any tissues they act in. Here *Lhx9* expression was not observed in the primordial follicles but was in all other stages of follicle growth. The reproductive stage at which this expression pattern begins is unknown. The earliest this has been characterised is five months. However, expression likely begins before this time, perhaps around puberty or oestrous onset, as is the case with several genes^35^.

There was also evidence of some expression in cells of the corpora lutea (Fig. 2). This may reflect expression in “small luteal cells” as described in the Mouse Cell Atlas single-cell expression data (Fig S1). These cells are not well characterised in terms of function compared to the better understood large luteal cells that are thought to be responsible for most of the progesterone expression^36,37^. Literature on their function does not offer many recent publications, but older literature highlights their thecal origin and role in progesterone synthesis and cholesterol transport^38^. The differentiation of the two populations is not well understood. The large luteal cells develop primarily from the granulosa cells, with evidence that they may also differentiate from the small luteal cells themselves^39^.

The RNA Sequencing analysis and subsequent GO analysis identified several differentially expressed genes and terms related to cholesterol metabolism and steroid synthesis (Fig. 6). These gene changes may reflect changes in the luteal cells expressing *Lhx9* described above. Large luteal cells have been shown to express steroidogenesis genes at greater levels than their small counterparts with elevated levels of cholesterol transport. If there is an altered differentiation of luteal cells in the *Lhx9*^+/-^ ovary, this may impact the expression of cholesterol and steroid-related genes. *Lhx9* expression in the steroidal cells of the male Leydig cells has been shown previously^40,41^, thus the expression of *Lhx9* shown here in the steroidal corporal lutea may reflect a function in the differentiation of steroidogenic cells. Genes with shared functionality in the testes and ovary include ephrins, these genes having a dual function in the Leydig and luteal/granulosa cells^42^. Ephrin proteins are also involved in epithelial proliferation, among other things^43^. This may be consistent with the function of *Lhx9* in regulating differentiation in embryonic tissues^44,45^.

### Impact of reduced Lhx9 expression on the ovary

Given that *Lhx9* is expressed in both the developing and adult ovaries, it is difficult to determine precisely at which point the reduced Lhx9 expression contributes to the subfertility observed in the *Lhx9*^+/-^ mouse line. Are the phenotypes in the adult a consequence of abnormal cellular dynamics during development? Or is it a result of functional changes occurring in the adult tissue due to aberrant gene regulation? To clarify this question, we carried out some fundamental histological analysis on the *Lhx9*^+/-^ ovaries to determine if there were any structural changes and RNA sequencing to assess the impact of reduced *Lhx9* on a transcriptional level.

The absence of any apparent changes in gross ovarian structure or follicular anatomy indicates that the haploinsufficiency of *Lhx9* is unlikely to result in any large-scale structural changes that arise during development. The presence of follicles at a range of growth stages and no observed abnormalities in the oestrous cycle also mean that the reduced *Lhx9* is unlikely to impact follicle growth and ovulation in the ovary. There were no differences in the number of antral and pre-antral follicles between the wildtype and heterozygous ovaries. Together these results show that the fertility decline of *Lhx9*^+/-^ mice are unlikely due to dysfunctional ovulation. The lack of difference observed in large follicle numbers means that analysing differences on a transcriptional level provided a more practical insight. The gene expression changes do not merely reflect differing follicle numbers or structural makeup. The number of significant differentially expressed genes is relatively small due to using a heterozygous mouse line instead of knockout mice. However, the considerable fold changes identified using only three replicates underline the regulatory role of *Lhx9* in the ovary.

### Lhx9 in steroidal cells of the ovary

GO analysis revealed that the differentially expressed genes are involved in significant processes and pathways vital to the functioning of an ovary. Downregulated biological processes, functional clusters, and pathway analysis highlight a downregulation of genes involved in steroid synthesis, cholesterol metabolism, and lipid metabolism. These processes are tightly intertwined, with the metabolising of lipids being crucial as a source of cholesterol and cholesterol being an essential precursor for many steroid hormones^46,47^. While these genes are vital for ovarian steroidogenesis, the metabolic pathways they are a part of consist of many genes. Thus, there is a level of compensation within the system. However, given the vital role ovarian steroids play in fertility, the changes in the expression of these genes are a likely source of the subfertility exhibited by the *Lhx9* mice, even if the exact mechanism of this is not yet known. Looking to studies utilising knockout mouse models for these genes can indicate the potential impact in the *Lhx9*^+/-^ model. The primary site of expression of 17β-Hydroxysteroid Dehydrogenase Type 7 (*Hsd17b7*) in the mouse is the large luteal cells of the ovary, but it is also expressed in the granulosa and theca cells^48^. It is part of a group of enzymes that convert keto-steroids to hydroxy-steroids and vice versa; in addition, *Hsd17b7* is involved in cholesterol metabolism^49^. As *Hsd17b7*^-/-^ mice are embryonic lethal, there is little indication of the impact of reduced expression in the adult on steroid expression. Cellular expression of High-Density Lipoprotein Receptor (*Srb1*) is found in the granulosa and theca cells of the ovary and interstitial tissue. *Srb1* deficient mice are sterile, with elevated plasma cholesterol and an inability to form multicell embryos^50^. Despite this, *Srb1*^-/-^ mice exhibit normal cyclicity and normal steroid levels. If the differentiation of luteal cells is impacted, this may cause the reduced *Hsd17b7* expression. Therefore, altered steroid synthesis and metabolism are a likely cause of the subfertility exhibited by *Lhx9*^+/-^ mice.

### Lhx9 in the ovarian surface epithelium

Expression in the ovarian surface epithelium and the luminal epithelium of the uterine tubes may indicate a tissue maintenance role for *Lhx9.* We know that the ovarian surface epithelium has a distinct population of multipotent cells that undergo regular expansion and differentiation to replace damaged cells lost during ovulation^51,52^. Bowen et al. determined that *Lhx9* was a critical stem cell marker expressed in the OSE that became downregulated in ovarian cancer epithelium. This implies that LHX9 may suppress the proliferation and differentiation of the epithelium instead of maintaining a stem-like state for the average turnover of the ovary epithelium. Read counts from the RNA-seq dataset in our study showed an increase in expression of epithelial marker gene *Prom1* in the *Lhx9*^+/-^ ovaries (Fig. 8), indicating that there may be an increase in epithelial cell proliferation in response to the reduced *Lhx9*. The GO analysis of significantly upregulated genes highlighted several epithelial related terms and a cancer-related pathway as being over-represented (Fig. 7). Several of the upregulated genes identified as part of these enriched GO terms show evidence for contribution to ovarian carcinoma. *Hoxa10* and *Cd24* are not expressed in normal OSE but are highly expressed in cancerous ovarian tissues^30,53,54^. In addition, analysis of transcription factor targets using the ChEA database determined that gene targets of SUZ12, JARID2, EZH2 and TP53 were significantly overrepresented in the list of genes upregulated in the *Lhx9*^+/-^ ovary. These genes have been characterised as contributing to ovarian carcinoma pathology^24,26,27,55^.

*LHX9* has been downregulated in other cancers, including cervical cancer and gliomas, where it acts to bind directly to p53^56^. The downregulation of *LHX9* gene expression is essentially methylation mediated, as shown by high levels of methylation in 88% of high-grade glioma field^57^ and liver^58^. In both OSE and other cancers, high expression levels of *Lhx9 are* correlated with a good prognosis^56^. Reduction of LHX9 promotes the expression of *PGK1*, a rate-limiting factor in the glycolysis process; glycolysis is a significantly upregulated process in many tumours, including gliomas^56^. In cervical cancer, *LHX9* has been proposed as a valuable biomarker for early detection^59^. Additionally, it has been shown that a knockdown of *LHX9* significantly impairs the proliferation of osteosarcoma cells^60^. In mice with a p53 point mutation (*Trp53^R172H^*), *Lhx9* expression was significantly reduced in a heterozygous mutant and undetectable in the ovarian cancer cells of a homozygous mutant^61^.

The uterine tube originates from the Müllerian duct that invaginates from the coelomic epithelium in the embryonic ovary. A lack of coelomic proliferation results in the gonadal agenesis typical of *Lhx9*^-/-^ mice^14^. The shared origin between the uterine tube and ovarian epithelia may account for the persistent expression of *Lhx9* and other shared markers such as *Lgr5*^62^. A more detailed analysis of LHX9 protein expression in the uterine epithelium was carried out as part of a study by Auersperg et al. in 2013. Expression of LHX9 was strongly detected in the nuclei of the fimbriae epithelium and all cell types, including ciliated cells and secretory cells^63^. Together this evidence points to a role for LHX9 in maintaining the appropriate level of turnover in the UTE and OSE, maintaining the stem population whilst controlling cellular differentiation. To investigate this further, cell tracing studies would be helpful to identify which cells LHX9+ epithelia contribute to in the adult ovary. Additionally, analysis of expression changes in the OSE in *Lhx9*^+/-^ mice could be further defined with single-cell RNA sequencing tools. A focused investigation into the impact of altered *Lhx9* expression on ovarian cancer cell lines will provide further insight into its role in healthy OSE and ovarian cancer progression.

## Conclusions

In this study, we have shown the expression of *Lhx9* in the somatic and germ cells of the adult mouse ovary. *Lhx9*^+/-^ mice exhibit reduced litters over time than their wildtype littermates. However, we have shown this is not due to any significant morphological changes in follicle structure or the primary function of the ovary. The transcriptional changes we observed following RNA-sequencing highlight the need for further investigation into the role of *Lhx9* in adult tissues like the ovary. Possible roles in the differentiation of somatic populations like luteal cells and the ovarian epithelia may have implications for fertility and cancer research.

## Methods

### Animal husbandry and tissue collection

To understand the role of *Lhx9* in the adult ovary, we used an *Lhx9* knockout mouse line obtained from the Jackson Laboratory (*Lhx9^tm1Lmgd^/EiJ*) and backcrossed onto a C57BL/6 strain at Otago. Heterozygous knockout *Lhx9*^+/-^ mice were used for the comparisons in this study. Mice were housed in standard conditions with *ad libitum* access to food and water. The oestrous cycle of mice was observed by vaginal lavage followed by staining with toluidine blue to observe vaginal cytology. Oestrous stages were classified according to well-established protocols ^64^. Mice were culled via cervical dislocation at five months of age during the proestrus stage of the oestrous cycle. Tissue was collected under sterile conditions and fixed in 4% PFA for sectioning or stored in RNALater (ThermoFisher) for RNA extraction. All animal procedures were carried out under the approval of the University of Otago Animal Ethics Committee.

### Genotyping

Tissue biopsies were first digested and then used for PCR amplification to determine wildtype (*Lhx9*^+/+^) or heterozygote (*Lhx9*^+/-^) status. Primers were designed to amplify the Neomycin cassette insert indicating a mutant allele. In addition, primers were designed to amplify the wildtype *Lhx9* region, acting as an internal control (Table S1).

### In situ hybridisation and Immunohistochemistry

Ovaries were embedded in either OCT or paraffin wax depending on the downstream application. Serial sections were cut at 5 μm thickness from wax and 10 μm thickness from frozen blocks. In situ hybridisation was carried out using probes designed to amplify all isoforms of the *Lhx9* mRNA^12^.Standard in situ protocols were carried out using frozen sections^65^. This consisted of tissue fixing in 4% PFA, followed by stringency washes and acetylation of tissue samples. Probes were allowed to bind overnight at 60°C. Following removal of the probe, the tissue was blocked against non-specific binding using heat-inactivated sheep serum and bovine serum albumin. Anti-DIG antibody (1 in 2000) was added, and slides were left overnight at 4°C. The stain was developed at room temperature following antibody removal, followed by stain removal and overnight fixing in 4% PFA. Slides were mounted in 70% glycerol.

Immunohistochemistry was carried out using paraffin-embedded sections. A standard laboratory protocol was optimised for use in the ovary^66^. This consisted of de-paraffinizing and rehydration washes in xylene/ethanol, followed by antigen retrieval in sodium citrate buffer for 30 min in a medium powered microwave. Sections were allowed to cool for 30 min, followed by washes to remove sodium citrate buffer. Blocking was performed for 1 h in a 10% heat-inactivated sheep serum solution and 1 % BSA diluted in 1% PBTx. Primary antibodies were diluted at 1 in 1000 in the blocking buffer and incubated overnight at 4°C. The primary antibody was removed using a series of washes in PBSx and a 5 min incubation in 3% H_2_O_2_. Blocking was carried out for 20 min in the same blocking buffer at room temperature. Secondary antibody (Abcam, ab205718) was diluted in blocking buffer at 1:2000 and added to sections for 2 h. Staining was carried out using the DAB detection kit (Abcam, ab64238). Slides were counterstained in 50% Haematoxylin and taken through a reverse ethanol/xylene series before mounting in DPX.

### Large follicle counting

Following dissection, they were fixed in 4% PFA to prepare ovaries for follicle counting. Ovaries were embedded in paraffin wax, stained under a standard haematoxylin and eosin protocol, and then sectioned serially at 5 μm thick. Each fifth section was used for counting, and follicles were classified according to the following criteria. Antral follicles were classed as having the presence of an antrum within the granulosa cell layer. Preantral follicles were classed as any follicle containing two or more layers of granulosa cells where no antrum was present. Pre-ovulatory follicles were classed as having a diameter greater than 250 μm. All counts were carried out blinded to the *Lhx9* genotype.

### RNA Extraction and Sequencing

RNA extraction was carried out using Trizol (Ambion), followed by chloroform extraction and ethanol precipitation. Samples were treated with Invitrogen Purelink DNase to remove any genomic DNA contamination and resuspended in MilliQ H_2_O. RNA concentration and quality were determined on a Nanodrop, and the RIN number was assessed using a BioAnalyzer. A total of three replicates per genotype were collected. Library preparation and sequencing were performed on an Illumina HiSeq platform generating 150 bp paired-end or single-end reads. Raw sequence data is available in the Sequence Read Archive under project: PRJNA816143.

Initial quality control, mapping and alignment were carried out on Galaxy^67^. The quality of the reads was determined by FastQC (v 0.11.8). The files were then mapped to the mouse genome (mm10) using HISAT2 (v 2.1.0). Read alignment was visualised on UCSC through Bam Coverage. featureCounts (v 1.6.4) was used to assign reads to genes, and from these files, a counts matrix was generated using Column Join (v 0.0.3).

Differential gene expression analysis was performed using R Studio with EdgeR ^68^ and DESeq2 ^69^. First, read counts were normalised using RUVSeq ^70^. Differentially expressed genes were considered significant when the False Discovery Rate (FDR) adjusted p-value was <0.05. DEG list is provided in supplementary File 1. Gene Ontology analysis was carried out using DAVID ^71^, KOBAS incorporating the KEGG and Panther databases ^72–74^, and ShinyGO, ^75^. A complete list of genes and their GO terms are found in supplementary File 2.

### RT-qPCR validation

The sequencing results were validated using RNA collected from the ovaries of different individuals from separate litters. RNA was reverse transcribed using iScript reverse transcriptase (Biorad). RT-qPCR was carried out using two ng of cDNA loaded in triplicate. RT-qPCR was performed using SYBR Green master mix (ThermoFisher) on a ViiA 7 PCR Machine (ThermoFisher). Primers were designed for a selection of the differentially expressed genes. Primer sequences can be found in Supplementary Table S1. Relative expression was measured against *Rps29* and *Actinb* reference genes using the 2^(CT^ref^-CT^target^) method.

### Statistical Analysis

All results are expressed as mean ±SEM. Analysis was carried out in Prism 7 software. An unpaired t-test determined statistical significance. Statistical significance labelled as * p <0.05, ** p <0.01, *** p <0.001, **** p <0.0001.

## Acknowledgements

We wish to thank Dr Michael Pankhurst for his advice and guidance on ovarian stereology. We also thank Berivan Temiz for her feedback on this manuscript. We thank the Department of Anatomy for funding. SW was supported by a University of Otago PhD scholarship.

## Author contributions

S.W carried out experimental work and wrote the manuscript. M.W assisted with experimental design and with manuscript revision.

