## Supplementary Information for "Lhx9 haploinsufficiency alters transcription in the adult mouse ovary, causing subfertility and abnormal epithelium"

**Supplementary Tables and Figures**

Table S1: Primers used for genotyping and RNA-Sequencing Validation (5'-3')

| Target | Forward Primer | Reverse Primer |
|---|---|---|
| *Lhx9* | GAATGTTGAAAATGGGGGGAA | TAAGATGCCACTGTTTGTCTACGG |
| *Neomycin* | TGCTCTTCGTCCAGATCA | CTGCCGAGAAAGTCTCCATC |
| *Snap25* | CCATCAGTGGTGGCTTCATC | CACCTGCTCCAGGTTCTCAT |
| *Claudin18* | GCTGGTGACCAACTTCTGGA | GAACAGAGCTGCACCAAAGG |
| *Hsd17b7* | CTGTGACACCGTACAACGGA | GCTCGGGTGATCCGATTTCT |
| *Tmprs11g* | CTCTGCGGTGCTTCTCTCAT | TGTCCACAGTTTGGGGTTTT |
| *Slc5a1* | CCCATGTTCCTCATGGTGAT | ACTGGTGTGCCGCAGTATTT |
| *Clca1* | AGCAGCACCTCCGAAGAAC | GCACAGCTGCTTGTCTTGAA |
| *Sprr2f* | GGTACACACGTCCTGGAATAC | CAGCACCTAGGAAGACCATAAA |
| *Rps29* | TGAAGGCAAGATGGGTCAC | GCACATGTTCAGCCCGTATT |

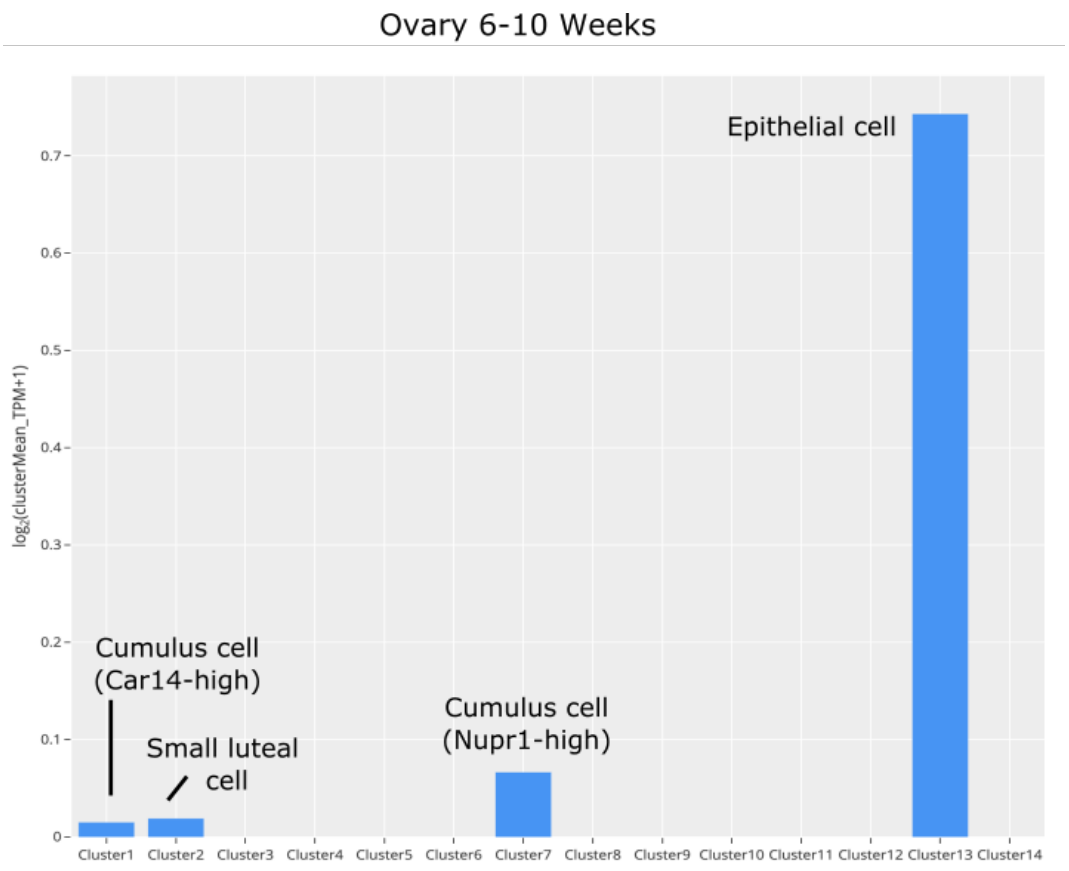


**Figure S1:** *Lhx9* expression in clusters identified by the Mouse Cell Atlas[254] in the adult ovary Expression shown as $\log_2$ (clusterMean_Transcript counts per million +1). *Lhx9* expression is present in four of 14 clusters of the adult mouse ovary (aged 6-10 weeks).
