## Supplementary material for "Lhx9 haploinsufficiency alters transcription in the adult mouse ovary, causing subfertility and abnormal epithelium": File S1

**Down in Het**

| Gene ID | Gene Name | Log2 Fold Change (Edge R) | p-adjusted | FDR |
|---|---|---|---|---|
| 11302 | Aatk | 1.67 | 3.19E-07 | 0.00018423 |
| 12032 | Bcan | 1.32 | 0.00038883 | 0.04276892 |
| 12393 | Runx2 | 1.21 | 0.00022222 | 0.02951211 |
| 12555 | Cdh15 | 1.40 | 0.00036796 | 0.0421622 |
| 14058 | F10 | 1.48 | 5.00E-05 | 0.01017752 |
| 15490 | Hsd17b7 | 2.32 | 5.51E-07 | 0.00028231 |
| 16493 | Kcna5 | 1.39 | 0.0004497 | 0.04687549 |
| 16891 | Lipg | 1.59 | 0.00077058 | 0.06639281 |
| 16979 | Lrrn1 | 1.29 | 4.71E-05 | 0.0099554 |
| 16987 | Lss | 1.33 | 0.00079245 | 0.06694522 |
| 20209 | Saa2 | 1.76 | 1.28E-05 | 0.00360919 |
| 20614 | Snap25 | 3.73 | 2.03E-09 | 5.21E-06 |
| 20778 | Scarb1 | 1.30 | 2.99E-05 | 0.00704544 |
| 20964 | Syn1 | 1.52 | 4.72E-05 | 0.0099554 |
| 23928 | Lamc3 | 1.16 | 0.0001368 | 0.02139442 |
| 57435 | Plin4 | 1.26 | 2.77E-05 | 0.00661044 |
| 66857 | Plbd1 | 1.01 | 0.00087259 | 0.07078998 |
| 67473 | Slc47a1 | 1.30 | 8.52E-05 | 0.01558302 |
| 73166 | Tm7sf2 | 1.11 | 0.00119437 | 0.08704788 |
| 83674 | Cnnm1 | 1.19 | 8.95E-05 | 0.01604967 |
| 215332 | Slc36a3 | 1.68 | 4.99E-05 | 0.01017752 |
| 223920 | Soat2 | 1.49 | 7.23E-05 | 0.01379957 |
| 243612 | Ssuh2 | 1.70 | 0.00019914 | 0.02762333 |
| 269615 | Plch2 | 1.22 | 6.72E-05 | 0.01310291 |
| 381399 | Bpifb4 | 2.02 | 0.00010846 | 0.01834463 |

**Up in Het**

| Gene ID | Gene Name | Log2 Fold Change (Edge R) | p-adjusted | FDR |
|---|---|---|---|---|
| 11658 | Alcam | -1.15 | 0.00038432 | 0.04276892 |
| 11833 | Aqp8 | -2.69 | 0.00000064 | 0.00031785 |
| 11846 | Arg1 | -2.18 | 0.00000006 | 0.00005370 |
| 12349 | Ca2 Car2 | -1.63 | 0.00003860 | 0.00854798 |
| 12484 | Cd24 Cd24a Ly-52 | -1.32 | 0.00002490 | 0.00620297 |
| 12491 | Cd36 | -0.98 | 0.00098895 | 0.07708590 |
| 12550 | Cdh1 | -1.21 | 0.00027341 | 0.03407066 |
| 12556 | Cdh16 | -2.75 | 0.00000427 | 0.00139159 |
| 12870 | Cp | -1.21 | 0.00068277 | 0.06277609 |
| 12931 | Crlf1 Crlm3 | -2.76 | 0.00000000 | 0.00000000 |
| 13082 | Cyp26a1 Cyp26 P450ra | -1.88 | 0.00000082 | 0.00038895 |
| 13175 | Dclk1 Dcamkl1 Dclk | -1.16 | 0.00106060 | 0.07998822 |
| 13723 | Emb Gp70 | -1.17 | 0.00016441 | 0.02374881 |
| 14013 | Mecom Evi1 Mds1 Prdm3 | -1.04 | 0.00096702 | 0.07571028 |
| 14348 | Fut9 | -1.77 | 0.00000273 | 0.00100000 |
| 14425 | Galnt3 | -1.51 | 0.00019037 | 0.02666528 |
| 14594 | Ggta1 Ggta-1 | -0.97 | 0.00099376 | 0.07708590 |
| 14619 | Gjb2 Cxn-26 | -2.73 | 0.00000010 | 0.00008360 |
| 15395 | Hoxa10 Hox-1.8 Hoxa-10 | -2.44 | 0.00000349 | 0.00119859 |
| 16616 | Klk1b21 Klk-21 Klk21 | -2.27 | 0.00000000 | 0.00000859 |
| 17022 | Lum Lcn Ldc | -0.99 | 0.00073917 | 0.06509374 |
| 17392 | Mmp3 | -2.80 | 0.00000029 | 0.00018016 |
| 17951 | Naip5 Birc1e Naip-rs3 | -1.63 | 0.00009620 | 0.01673784 |
| 18300 | Fam3d Oit1 | -1.96 | 0.00000036 | 0.00019667 |
| 18546 | Pcp4 Pep19 | -2.74 | 0.00000316 | 0.00111205 |
| 18599 | Padi1 Pad1 Pdi1 | -2.10 | 0.00000014 | 0.00010408 |
| 18600 | Padi2 Pad2 Pdi Pdi2 | -2.33 | 0.00000006 | 0.00005370 |
| 18667 | Pgr Nr3c3 Pr | -1.65 | 0.00015911 | 0.02323843 |
| 18703 | Pigr | -2.40 | 0.00000002 | 0.00002340 |
| 19126 | Prom1 Prom Proml1 | -1.34 | 0.00038601 | 0.04276892 |
| 19225 | Ptgs2 Cox-2 Cox2 Pghs-b Tis10 | -1.83 | 0.00000803 | 0.00244018 |
| 20192 | Ryr3 | -1.54 | 0.00001680 | 0.00455252 |
| 20390 | Sftpd Sftp4 | -1.72 | 0.00015042 | 0.02228769 |
| 20522 | Slc23a1 Svct1 Yspl3 | -1.19 | 0.00017453 | 0.02483430 |
| 20531 | Slc34a2 Npt2b | -1.22 | 0.00056065 | 0.05462980 |
| 20538 | Slc6a2 | -2.82 | 0.00000002 | 0.00002060 |
| 20753 | Sprr1a | -3.95 | 0.00000223 | 0.00088941 |
| 22095 | Tshr | -1.39 | 0.00086710 | 0.07066429 |
| 23795 | Agr2 Gob4 | -1.64 | 0.00009200 | 0.01619672 |
| 23844 | Clca1 Clca3 Gob5 | -6.38 | 0.00000000 | 0.00000000 |
| 50528 | Tmprss2 | -1.53 | 0.00007590 | 0.01418055 |
| 50706 | Postn Osf2 | -2.02 | 0.00000176 | 0.00074945 |
| 53315 | Sult1d1 St1d1 | -3.37 | 0.00000262 | 0.00097701 |
| 54486 | Hpgds Gsts Pgds Ptgds2 | -1.30 | 0.00014439 | 0.02192721 |
| 56429 | Dpt | -1.56 | 0.00042326 | 0.04501501 |
| 56753 | Tacstd2 Trop2 | -2.37 | 0.00000423 | 0.00139159 |
| 58222 | Rab37 | -1.85 | 0.00000016 | 0.00011658 |
| 66889 | Rnf128 Grail Greul1 MNCb-3816 | -1.37 | 0.00065447 | 0.06079749 |
| 70405 | Calml3 | -1.97 | 0.00000030 | 0.00018016 |
| 71091 | Cdkl1 | -1.22 | 0.00011192 | 0.01875406 |
| 74145 | F13a1 F13a | -1.42 | 0.00012239 | 0.02013155 |
| 76459 | Ca12 Car12 | -1.55 | 0.00012199 | 0.02013155 |
| 76477 | Pcolce2 Pcpe2 | -1.99 | 0.00000911 | 0.00272211 |
| 77522 | Tmem213 | -2.79 | 0.00000000 | 0.00000281 |
| 100689 | Spon2 | -1.75 | 0.00000354 | 0.00119859 |
| 114304 | Slc28a3 Cnt3 | -1.73 | 0.00002530 | 0.00620297 |
| 209558 | Enpp3 | -1.73 | 0.00000650 | 0.00204337 |
| 216749 | Nmur2 | -1.87 | 0.00000092 | 0.00041074 |
| 224116 | Muc20 | -1.92 | 0.00001290 | 0.00360919 |
| 232983 | Cxcl17 Vcc1 | -2.63 | 0.00000000 | 0.00000180 |
| 242125 | Mab21L3 | -1.85 | 0.00000183 | 0.00076381 |
| 242721 | Klhdc7a | -1.03 | 0.00074065 | 0.06509374 |
| 269643 | Ppp2r2c | -1.44 | 0.00013723 | 0.02139442 |
| 319767 | Atp10b | -1.33 | 0.00046461 | 0.04759975 |
| 394432 | Ugt1a7 Ugt1a7c | -1.26 | 0.00004850 | 0.01010769 |
| 394436 | Ugt1a1 Ugt1 | -1.42 | 0.00001070 | 0.00309762 |
