## Supplementary material for "Lhx9 haploinsufficiency alters transcription in the adult mouse ovary, causing subfertility and abnormal epithelium": File S2

**DAVID terms Downregulated  (lower expression in Het)**

| **Term BP** | Count | % | PValue | Genes | List Total | Pop Hits | Pop Total | Fold Enrichment |
|---|---|---|---|---|---|---|---|---|
| GO:0006629~lipid metabolic process | 7 | 28 | 1.02E-05 | 269615, 16891, 66857, 16987, 73166, 15490, 223920 | 22 | 459 | 18082 | 12.53 |
| GO:0006695~cholesterol biosynthetic process | 3 | 12 | 6.24E-04 | 16987, 73166, 15490 | 22 | 32 | 18082 | 77.05 |
| GO:0006694~steroid biosynthetic process | 3 | 12 | 0.0023 | 16987, 73166, 15490 | 22 | 62 | 18082 | 39.77 |
| GO:0042632~cholesterol homeostasis | 3 | 12 | 0.0024 | 16891, 20778, 223920 | 22 | 63 | 18082 | 39.14 |
| GO:0016042~lipid catabolic process | 3 | 12 | 0.0070 | 269615, 16891, 66857 | 22 | 109 | 18082 | 22.62 |
| GO:0034375~high-density lipoprotein particle remodeling | 2 | 8 | 0.0139 | 16891, 20778 | 22 | 12 | 18082 | 136.98 |
| GO:0043691~reverse cholesterol transport | 2 | 8 | 0.0150 | 16891, 20778 | 22 | 13 | 18082 | 126.45 |
| GO:0007269~neurotransmitter secretion | 2 | 8 | 0.0320 | 20614, 20964 | 22 | 28 | 18082 | 58.71 |
| GO:0033344~cholesterol efflux | 2 | 8 | 0.0320 | 20778, 223920 | 22 | 28 | 18082 | 58.71 |

| **Term CC** | Count | % | PValue | Genes | List Total | Pop Hits | Pop Total | Fold Enrichment |
|---|---|---|---|---|---|---|---|---|
| GO:0005783~endoplasmic reticulum | 7 | 28 | 0.004 | 16493, 16987, 14058, 73166, 15490, 11302, 223920 | 25 | 1323 | 19662 | 4.16 |
| GO:0016020~membrane | 16 | 64 | 0.007 | 269615, 215332, 16493, 73166, 15490, 20614, 20778, 223920, 57435, 83674, 16987, 14058, 11302, 67473, 16979, 12555 | 25 | 6998 | 19662 | 1.80 |
| GO:0043231~intracellular membrane-bounded organelle | 5 | 20 | 0.012 | 57435, 16987, 14058, 73166, 20778 | 25 | 751 | 19662 | 5.24 |
| GO:0043229~intracellular organelle | 2 | 8 | 0.016 | 20614, 20964 | 25 | 13 | 19662 | 121.00 |
| GO:0044295~axonal growth cone | 2 | 8 | 0.034 | 20614, 11302 | 25 | 28 | 19662 | 56.18 |

| **Term MF** | Count | % | PValue | Genes | List Total | Pop Hits | Pop Total | Fold Enrichment |
|---|---|---|---|---|---|---|---|---|
| GO:0005509~calcium ion binding | 4 | 16 | 0.044 | 269615, 12032, 14058, 12555 | 21 | 699 | 17446 | 4.75 |

**Cluster analysis**

| **Enrichment Score: 3.18** | **Annotation Cluster 1** | | | | |
|---|---|---|---|---|---|
| Term | Category | Count | % | Genes | PValue |
| mmu00100:Steroid biosynthesis | KEGG_PATHWAY | 4 | 16 | 16987, 73166, 15490, 223920 | 2.08E-06 |
| Lipid metabolism | UP_KEYWORDS | 7 | 28 | 269615, 16891, 66857, 16987, 73166, 15490, 223920 | 3.79E-06 |
| GO:0006629~lipid metabolic process | GOTERM_BP_DIRECT | 7 | 28 | 269615, 16891, 66857, 16987, 73166, 15490, 223920 | 1.02E-05 |
| GO:0006695~cholesterol biosynthetic process | GOTERM_BP_DIRECT | 3 | 12 | 16987, 73166, 15490 | 6.24E-04 |
| Steroid biosynthesis | UP_KEYWORDS | 3 | 12 | 16987, 73166, 15490 | 7.37E-04 |
| GO:0006694~steroid biosynthetic process | GOTERM_BP_DIRECT | 3 | 12 | 16987, 73166, 15490 | 0.0023 |
| GO:0005783~endoplasmic reticulum | GOTERM_CC_DIRECT | 7 | 28 | 16493, 16987, 14058, 73166, 15490, 11302, 223920 | 0.0043 |
| Lipid biosynthesis | UP_KEYWORDS | 3 | 12 | 16987, 73166, 15490 | 0.0123 |
| mmu01130:Biosynthesis of antibiotics | KEGG_PATHWAY | 3 | 12 | 16987, 73166, 15490 | 0.0359 |

| **Enrichment Score: 2.78** | **Annotation Cluster 2** | | | | |
|---|---|---|---|---|---|
| Term | Category | Count | % | Genes | PValue |
| mmu00100:Steroid biosynthesis | KEGG_PATHWAY | 4 | 16 | 16987, 73166, 15490, 223920 | 2.08E-06 |
| Lipid metabolism | UP_KEYWORDS | 7 | 28 | 269615, 16891, 66857, 16987, 73166, 15490, 223920 | 3.79E-06 |
| GO:0006629~lipid metabolic process | GOTERM_BP_DIRECT | 7 | 28 | 269615, 16891, 66857, 16987, 73166, 15490, 223920 | 1.02E-05 |
| Lipid degradation | UP_KEYWORDS | 3 | 12 | 269615, 16891, 66857 | 0.0050 |
| GO:0016042~lipid catabolic process | GOTERM_BP_DIRECT | 3 | 12 | 269615, 16891, 66857 | 0.0070 |

| **Enrichment Score: 1.44** | **Annotation Cluster 3** | | | | |
|---|---|---|---|---|---|
| Term | Category | Count | % | Genes | PValue |
| Glycoprotein | UP_KEYWORDS | 12 | 48 | 16891, 16493, 381399, 23928, 12032, 66857, 14058, 15490, 16979, 20778, 20964, 12555 | 8.74E-04 |
| SM00181:EGF | SMART | 3 | 12 | 23928, 12032, 14058 | 0.012 |
| Secreted | UP_KEYWORDS | 6 | 24 | 16891, 381399, 23928, 12032, 20209, 14058 | 0.029 |
| IPR000742:Epidermal growth factor-like domain | INTERPRO | 3 | 12 | 23928, 12032, 14058 | 0.031 |
| glycosylation site:N-linked (GlcNAc...) | UP_SEQ_FEATURE | 10 | 40 | 16891, 16493, 381399, 23928, 12032, 14058, 15490, 16979, 20778, 12555 | 0.034 |
| Signal | UP_KEYWORDS | 10 | 40 | 16891, 381399, 23928, 12032, 66857, 20209, 14058, 11302, 16979, 12555 | 0.036 |
| Disulfide bond | UP_KEYWORDS | 8 | 32 | 16891, 381399, 23928, 12032, 66857, 14058, 16979, 20778 | 0.038 |
| signal peptide | UP_SEQ_FEATURE | 9 | 36 | 16891, 381399, 23928, 12032, 66857, 20209, 14058, 16979, 12555 | 0.044 |

| **Enrichment Score: 1.18** | **Annotation Cluster 4** | | | | |
|---|---|---|---|---|---|
| Term | Category | Count | % | Genes | PValue |
| Cell membrane | UP_KEYWORDS | 9 | 36 | 57435, 269615, 16493, 83674, 15490, 20614, 67473, 20778, 12555 | 0.034 |
| Membrane | UP_KEYWORDS | 15 | 60 | 269615, 215332, 16493, 73166, 15490, 20614, 20778, 223920, 57435, 83674, 16987, 11302, 67473, 16979, 12555 | 0.037 |

| **Enrichment Score: 1.03** | **Annotation Cluster 5** | | | | |
|---|---|---|---|---|---|
| Term | Category | Count | % | Genes | PValue |
| GO:0016020~membrane | GOTERM_CC_DIRECT | 16 | 64 | 269615, 215332, 16493, 73166, 15490, 20614, 20778, 223920, 57435, 83674, 16987, 14058, 11302, 67473, 16979, 12555 | 0.00653 |
| Membrane | UP_KEYWORDS | 15 | 60 | 269615, 215332, 16493, 73166, 15490, 20614, 20778, 223920, 57435, 83674, 16987, 11302, 67473, 16979, 12555 | 0.03676 |
| tran+B26:B66smembrane region | UP_SEQ_FEATURE | 11 | 44 | 215332, 16493, 83674, 73166, 15490, 11302, 67473, 16979, 20778, 223920, 12555 | 0.04169 |

**DAVID terms Upregulated (higher expression in het)**

| **Term BP** | Count | % | PValue | Genes | List Total | Pop Hits | Pop Total | Fold Enrichment |
|---|---|---|---|---|---|---|---|---|
| GO:0007155~cell adhesion | 9 | 13.64 | 1.90E-04 | 11658, 12491, 50706, 12484, 12550, 13723, 100689, 56429, 12556 | 61 | 485 | 18082 | 5.50 |
| GO:0046697~decidualization | 3 | 4.55 | 0.003 | 19225, 14619, 12550 | 61 | 24 | 18082 | 37.05 |
| GO:0072139~glomerular parietal epithelial cell differentiation | 2 | 3.03 | 0.007 | 19126, 12484 | 61 | 2 | 18082 | 296.43 |
| GO:2000768~positive regulation of nephron tubule epithelial cell differentiation | 2 | 3.03 | 0.010 | 19126, 12484 | 61 | 3 | 18082 | 197.62 |
| GO:0007204~positive regulation of cytosolic calcium ion concentration | 4 | 6.06 | 0.013 | 12491, 12484, 232983, 216749 | 61 | 148 | 18082 | 8.01 |
| GO:0008228~opsonization | 2 | 3.03 | 0.016 | 20390, 100689 | 61 | 5 | 18082 | 118.57 |
| GO:0018101~protein citrullination | 2 | 3.03 | 0.016 | 18599, 18600 | 61 | 5 | 18082 | 118.57 |
| GO:0090336~positive regulation of brown fat cell differentiation | 2 | 3.03 | 0.020 | 19225, 14013 | 61 | 6 | 18082 | 98.81 |
| GO:0072112~glomerular visceral epithelial cell | 2 | 3.03 | 0.023 | 19126, 12484 | 61 | 7 | 18082 | 84.69 |
| GO:0043627~response to estrogen | 3 | 4.55 | 0.026 | 20531, 12484, 12349 | 61 | 75 | 18082 | 11.86 |
| GO:0042493~response to drug | 5 | 7.58 | 0.026 | 19225, 11846, 12550, 394436, 20538 | 61 | 339 | 18082 | 4.37 |
| GO:0015670~carbon dioxide transport | 2 | 3.03 | 0.026 | 76459, 12349 | 61 | 8 | 18082 | 74.11 |
| GO:0060907~positive regulation of macrophage cytokine production | 2 | 3.03 | 0.026 | 12491, 100689 | 61 | 8 | 18082 | 74.11 |
| GO:0010042~response to manganese ion | 2 | 3.03 | 0.026 | 19225, 11846 | 61 | 8 | 18082 | 74.11 |
| GO:0052697~xenobiotic glucuronidation | 2 | 3.03 | 0.029 | 394432, 394436 | 61 | 9 | 18082 | 65.87 |
| GO:0071318~cellular response to ATP | 2 | 3.03 | 0.039 | 19225, 20192 | 61 | 12 | 18082 | 49.40 |
| GO:1900025~negative regulation of substrate adhesion-dependent cell spreading | 2 | 3.03 | 0.039 | 56753, 50706 | 61 | 12 | 18082 | 49.40 |
| GO:0032355~response to estradiol | 3 | 4.55 | 0.044 | 19225, 14619, 50706 | 61 | 101 | 18082 | 8.80 |
| GO:0006693~prostaglandin metabolic process | 2 | 3.03 | 0.049 | 54486, 19225 | 61 | 15 | 18082 | 39.52 |
| GO:0071498~cellular response to fluid shear stress | 2 | 3.03 | 0.049 | 19225, 12349 | 61 | 15 | 18082 | 39.52 |

| **Term CC** | Count | % | PValue | Genes | List Total | Pop Hits | Pop Total | Fold Enrichment |
|---|---|---|---|---|---|---|---|---|
| GO:0005576~extracellular region | 18 | 27.27 | 0.00003 | 18300, 50528, 50706, 17022, 224116, 74145, 17392, 23795, 209558, 23844, 76477, 12870, 20390, 232983, 12931, 56429, 100689, 18703 | 65 | 1753 | 19662 | 3.106 |
| GO:0031528~microvillus membrane | 4 | 6.06 | 0.00005 | 19126, 20531, 12484, 224116 | 65 | 22 | 19662 | 54.999 |
| GO:0070062~extracellular exosome | 22 | 33.33 | 0.00007 | 20522, 19126, 50528, 18599, 18600, 12550, 17022, 209558, 11658, 56753, 71091, 11846, 76477, 12870, 70405, 16616, 12349, 56429, 100689, 14425, 12556, 18703 | 65 | 2674 | 19662 | 2.489 |
| GO:0005615~extracellular space | 16 | 24.24 | 0.00007 | 19126, 12491, 50706, 17022, 17392, 23795, 23844, 56753, 11846, 12870, 20390, 16616, 12349, 56429, 100689, 18703 | 65 | 1504 | 19662 | 3.218 |
| GO:0043231~intracellular membrane-bounded organelle | 9 | 13.636 | 0.003 | 19126, 19225, 11833, 12491, 13082, 14013, 394432, 20192, 394436 | 65 | 751 | 19662 | 3.625 |
| GO:0005578~proteinaceous extracellular matrix | 6 | 9.091 | 0.004 | 50706, 17022, 17392, 20390, 100689, 56429 | 65 | 316 | 19662 | 5.744 |
| GO:0045177~apical part of cell | 4 | 6.061 | 0.006 | 11833, 12491, 12550, 12349 | 65 | 115 | 19662 | 10.521 |
| GO:0009925~basal plasma membrane | 3 | 4.545 | 0.008 | 56753, 20522, 224116 | 65 | 42 | 19662 | 21.607 |
| GO:0005783~endoplasmic reticulum | 11 | 16.667 | 0.010 | 19126, 19225, 12491, 13082, 394432, 319767, 23795, 20192, 394436, 66889, 18667 | 65 | 1323 | 19662 | 2.515 |
| GO:0043005~neuron projection | 6 | 9.091 | 0.012 | 19225, 11846, 17951, 20538, 18546, 18667 | 65 | 420 | 19662 | 4.321 |
| GO:0016020~membrane | 33 | 50.000 | 0.012 | 19225, 242721, 19126, 12491, 50528, 13082, 20531, 12550, 14594, 394432, 224116, 114304, 394436, 20538, 56753, 14619, 22095, 20192, 216749, 13723, 12556, 20522, 12484, 66889, 209558, 11658, 11833, 77522, 76459, 12349, 14348, 14425, 18703 | 65 | 6998 | 19662 | 1.426 |
| GO:0016328~lateral plasma membrane | 3 | 4.545 | 0.013 | 56753, 14619, 12550 | 65 | 54 | 19662 | 16.805 |
| GO:0016021~integral component of membrane | 32 | 48.485 | 0.018 | 242721, 19126, 12491, 50528, 20531, 12550, 14594, 394432, 114304, 394436, 20538, 23844, 56753, 14619, 22095, 12870, 20192, 216749, 13723, 12556, 20522, 319767, 66889, 209558, 11658, 11833, 77522, 76459, 18667, 14348, 14425, 18703 | 65 | 6878 | 19662 | 1.407 |
| GO:0032580~Golgi cisterna membrane | 3 | 4.545 | 0.022 | 14594, 14425, 14348 | 65 | 70 | 19662 | 12.964 |
| GO:0016324~apical plasma membrane | 5 | 7.576 | 0.022 | 19126, 20522, 12491, 20531, 224116 | 65 | 328 | 19662 | 4.611 |
| GO:0005902~microvillus | 3 | 4.545 | 0.025 | 19126, 12349, 23844 | 65 | 76 | 19662 | 11.940 |
| GO:0005903~brush border | 3 | 4.545 | 0.026 | 19126, 20522, 20531 | 65 | 77 | 19662 | 11.785 |
| GO:0005887~integral component of plasma membrane | 9 | 13.636 | 0.029 | 56753, 19126, 11658, 11833, 22095, 114304, 394436, 20538, 13723 | 65 | 1126 | 19662 | 2.418 |
| GO:0030424~axon | 5 | 7.576 | 0.032 | 11658, 12550, 18546, 18667, 12349 | 65 | 370 | 19662 | 4.088 |

| **Term MF** | Count | % | PValue | Genes | List Total | Pop Hits | Pop Total | Fold Enrichment |
|---|---|---|---|---|---|---|---|---|
| GO:0046872~metal ion binding | 22 | 33.33 | 0.002 | 19225, 13082, 50706, 17951, 12550, 14013, 14594, 74145, 17392, 66889, 20538, 209558, 23844, 11846, 76459, 12870, 18667, 70405, 12349, 100689, 14425, 12556 | 60 | 3355 | 17446 | 1.91 |
| GO:0005509~calcium ion binding | 9 | 13.64 | 0.002 | 54486, 18599, 18600, 12550, 17392, 20192, 70405, 14425, 12556 | 60 | 699 | 17446 | 3.74 |
| GO:0016758~transferase activity, transferring hexosyl groups | 3 | 4.55 | 0.005 | 14594, 394432, 394436 | 60 | 31 | 17446 | 28.14 |
| GO:0016757~transferase activity, transferring glycosyl groups | 5 | 7.58 | 0.005 | 14594, 394432, 394436, 14425, 14348 | 60 | 208 | 17446 | 6.99 |
| GO:0004668~protein-arginine deiminase activity | 2 | 3.03 | 0.017 | 18599, 18600 | 60 | 5 | 17446 | 116.31 |
| GO:0001972~retinoic acid binding | 2 | 3.03 | 0.043 | 13082, 394436 | 60 | 13 | 17446 | 44.73 |
| GO:0004089~carbonate dehydratase activity | 2 | 3.03 | 0.050 | 76459, 12349 | 60 | 15 | 17446 | 38.77 |

**Cluster Analysis**

**Enrichment Score: 2.28** **Annotation Cluster 1**

| Term | Category | Count | % | Genes | PValue | List Total | Pop Hits | Pop Total |
|---|---|---|---|---|---|---|---|---|
| Extracellular matrix | UP_KEYWORDS | 6 | 9.09090909 | 50706, 17022, 17392, 20390, 100689, 56429 | 5.70E-04 | 66 | 235 | 22680 |
| GO:0005578~proteinaceous extracellular matrix | GOTERM_CC_DIRECT | 6 | 9.09090909 | 50706, 17022, 17392, 20390, 100689, 56429 | 0.0037 | 65 | 316 | 19662 |
| GO:0031012~extracellular matrix | GOTERM_CC_DIRECT | 4 | 6.06060606 | 50706, 17022, 17392, 56429 | 0.0709 | 65 | 294 | 19662 |

**Enrichment Score: 2.20** **Annotation Cluster 2**

| Term | Category | Count | % | Genes | PValue | List Total | Pop Hits | Pop Total |
|---|---|---|---|---|---|---|---|---|
| topological domain:Cytoplasmic | UP_SEQ_FEATURE | 22 | 33.33 | 19126, 50528, 12491, 20531, 12550, 14594, 114304, 20538, 209558, 11658, 56753, 11833, 14619, 22095, 76459, 216749, 18546, 13723, 14348, 14425, 12556, 18703 | 4.12E-04 | 63 | 2880 | 18012 |
| topological domain:Extracellular | UP_SEQ_FEATURE | 19 | 28.79 | 19126, 50528, 12491, 20531, 12550, 114304, 20538, 209558, 11658, 56753, 11833, 14619, 22095, 76459, 216749, 18546, 13723, 12556, 18703 | 4.24E-04 | 63 | 2256 | 18012 |
| transmembrane region | UP_SEQ_FEATURE | 28 | 42.42 | 242721, 19126, 12491, 50528, 20531, 12550, 14594, 394432, 114304, 394436, 20538, 56753, 14619, 22095, 216749, 13723, 12556, 20522, 66889, 209558, 11658, 11833, 77522, 76459, 18546, 14348, 14425, 18703 | 5.26E-04 | 63 | 4312 | 18012 |
| Transmembrane helix | UP_KEYWORDS | 31 | 46.97 | 242721, 19126, 12491, 50528, 20531, 12550, 14594, 394432, 114304, 394436, 20538, 56753, 14619, 22095, 12870, 20192, 216749, 13723, 12556, 20522, 319767, 66889, 209558, 11658, 11833, 77522, 76459, 18667, 14348, 14425, 18703 | 0.0059 | 66 | 6938 | 22680 |
| Transmembrane | UP_KEYWORDS | 31 | 46.97 | 242721, 19126, 12491, 50528, 20531, 12550, 14594, 394432, 114304, 394436, 20538, 56753, 14619, 22095, 12870, 20192, 216749, 13723, 12556, 20522, 319767, 66889, 209558, 11658, 11833, 77522, 76459, 18667, 14348, 14425, 18703 | 0.0061 | 66 | 6955 | 22680 |
| Membrane | UP_KEYWORDS | 36 | 54.55 | 19225, 242721, 19126, 12491, 50528, 13082, 20531, 12550, 14594, 394432, 224116, 114304, 394436, 20538, 56753, 14619, 22095, 12870, 20192, 216749, 13723, 12556, 20522, 12484, 319767, 66889, 209558, 11658, 11833, 77522, 76459, 18667, 12349, 14348, 14425, 18703 | 0.0077 | 66 | 8683 | 22680 |
| GO:0016020~membrane | GOTERM_CC_DIRECT | 33 | 50.00 | 19225, 242721, 19126, 12491, 50528, 13082, 20531, 12550, 14594, 394432, 224116, 114304, 394436, 20538, 56753, 14619, 22095, 20192, 216749, 13723, 12556, 20522, 12484, 66889, 209558, 11658, 11833, 77522, 76459, 12349, 14348, 14425, 18703 | 0.0125 | 65 | 6998 | 19662 |
| GO:0016021~integral component of membrane | GOTERM_CC_DIRECT | 32 | 48.48 | 242721, 19126, 12491, 50528, 20531, 12550, 14594, 394432, 114304, 394436, 20538, 23844, 56753, 14619, 22095, 12870, 20192, 216749, 13723, 12556, 20522, 319767, 66889, 209558, 11658, 11833, 77522, 76459, 18667, 14348, 14425, 18703 | 0.0181 | 65 | 6878 | 19662 |

\

**Enrichment Score: 1.65** **Annotation Cluster 3**

| Term | Category | Count | % | Genes | PValue | List Total | Pop Hits | Pop Total |
|---|---|---|---|---|---|---|---|---|
| Microsome | UP_KEYWORDS | 5 | 7.58 | 19225, 13082, 394432, 20192, 394436 | 5.64E-04 | 66 | 132 | 22680 |
| GO:0043231~intracellular membrane-bounded organelle | GOTERM_CC_DIRECT | 9 | 13.64 | 19126, 19225, 11833, 12491, 13082, 14013, 394432, 20192, 394436 | 0.0029 | 65 | 751 | 19662 |
| GO:0005783~endoplasmic reticulum | GOTERM_CC_DIRECT | 11 | 16.67 | 19126, 19225, 12491, 13082, 394432, 319767, 23795, 20192, 394436, 66889, 18667 | 0.0100 | 65 | 1323 | 19662 |

**Enrichment Score: 1.55** **Annotation Cluster 4**

| Term | Category | Count | % | Genes | PValue | List Total | Pop Hits | Pop Total |
|---|---|---|---|---|---|---|---|---|
| domain:Ig-like V-type 2 | UP_SEQ_FEATURE | 4 | 6.06 | 11658, 18546, 13723, 18703 | 4.25E-05 | 63 | 20 | 18012 |
| domain:Ig-like V-type 1 | UP_SEQ_FEATURE | 4 | 6.06 | 11658, 18546, 13723, 18703 | 4.94E-05 | 63 | 21 | 18012 |

**Enrichment Score: 1.45** **Annotation Cluster 5**

| Term | Category | Count | % | Genes | PValue | List Total | Pop Hits | Pop Total |
|---|---|---|---|---|---|---|---|---|
| Glycosyltransferase | UP_KEYWORDS | 5 | 7.58 | 14594, 394432, 394436, 14425, 14348 | 0.004 | 66 | 225 | 22680 |
| GO:0016758~transferase activity, transferring hexosyl groups | GOTERM_MF_DIRECT | 3 | 4.55 | 14594, 394432, 394436 | 0.005 | 60 | 31 | 17446 |
| GO:0016757~transferase activity, transferring glycosyl groups | GOTERM_MF_DIRECT | 5 | 7.58 | 14594, 394432, 394436, 14425, 14348 | 0.005 | 60 | 208 | 17446 |
| GO:0032580~Golgi cisterna membrane | GOTERM_CC_DIRECT | 3 | 4.55 | 14594, 14425, 14348 | 0.022 | 65 | 70 | 19662 |
| Signal-anchor | UP_KEYWORDS | 5 | 7.58 | 50528, 14594, 209558, 14425, 14348 | 0.037 | 66 | 439 | 22680 |
| Transferase | UP_KEYWORDS | 10 | 15.15 | 71091, 54486, 14594, 394432, 13175, 74145, 394436, 53315, 14425, | 0.045 | 66 | 1654 | 22680 |

**Enrichment Score: 1.33** **Annotation Cluster 6**

| Term | Category | Count | % | Genes | PValue | List Total | Pop Hits | Pop Total |
|---|---|---|---|---|---|---|---|---|
| Symport | UP_KEYWORDS | 3 | 4.55 | 20522, 20531, 20538 | 0.040 | 66 | 111 | 22680 |
| Sodium | UP_KEYWORDS | 3 | 4.55 | 20522, 20531, 20538 | 0.047 | 66 | 120 | 22680 |

**KOBAS Downregulated lower expression in HET)**

| Term | Database | ID | Input_number | Background number | P-Value | Pad | Input |
|---|---|---|---|---|---|---|---|
| Steroid biosynthesis | KEGG PATHWAY | mmu00100 | 4 | 19 | 1.23E-09 | 6.00E-07 | 223920\|16987\|15490\|73166 |
| Cholesterol metabolism | KEGG PATHWAY | mmu04979 | 3 | 49 | 5.28E-06 | 0.00043019 | 16891\|223920\|20778 |
| Androgen/estrogene/progesterone biosynthesis | PANTHER | P02727 | 2 | 13 | 4.25E-05 | 0.00148539 | 223920\|15490 |
| Synaptic vesicle trafficking | PANTHER | P05734 | 2 | 24 | 0.00013105 | 0.00291295 | 20964\|20614 |
| Ovarian steroidogenesis | KEGG PATHWAY | mmu04913 | 2 | 57 | 0.00068092 | 0.01040528 | 15490\|20778 |
| Metabolic pathways | KEGG PATHWAY | mmu01100 | 5 | 1494 | 0.0024933 | 0.02901129 | 16891\|16987\|15490\|73166\|269615 |
| Cholesterol biosynthesis | PANTHER | P00014 | 1 | 13 | 0.00908 | 0.03389404 | 16987 |

**KOBAS Upregulated (Higher expression in het)**

| Term | Database | ID | Input_number | Background number | P-Value | Pad | Input |
|---|---|---|---|---|---|---|---|
| Ascorbate and aldarate metabolism | KEGG PATHWAY | mmu00053 | 2 | 27 | 0.00114453 | 0.01648116 | 394436\|394432 |
| Glycosphingolipid biosynthesis - lacto and neolacto series | KEGG PATHWAY | mmu00601 | 2 | 27 | 0.00114453 | 0.01648116 | 14594\|14348 |
| Retinol metabolism | KEGG PATHWAY | mmu00830 | 3 | 91 | 0.00057994 | 0.01043899 | 394436\|394432\|13082 |
| Nitrogen metabolism | KEGG PATHWAY | mmu00910 | 2 | 17 | 0.00048742 | 0.01002702 | 12349\|76459 |
| Transcriptional misregulation in cancer | KEGG PATHWAY | mmu05202 | 4 | 183 | 0.00030911 | 0.00741865 | 50528\|17392\|15395\|19126 |
| Drug metabolism - cytochrome P450 | KEGG PATHWAY | mmu00982 | 3 | 68 | 0.00025434 | 0.00732511 | 394436\|394432\|54486 |
| Metabolism of xenobiotics by cytochrome P450 | KEGG PATHWAY | mmu00980 | 3 | 66 | 0.00023373 | 0.00732511 | 394436\|394432\|54486 |
| Porphyrin and chlorophyll metabolism | KEGG PATHWAY | mmu00860 | 3 | 41 | 6.09E-05 | 0.00292419 | 394436\|394432\|12870 |
| Metabolic pathways | KEGG PATHWAY | mmu01100 | 11 | 1494 | 4.75E-05 | 0.00292419 | 14425\|209558\|54486\|76459\|14348\|12349\|394436\|19225\|394432\|11846\|13082 |
| Chemical carcinogenesis | KEGG PATHWAY | mmu05204 | 4 | 94 | 2.54E-05 | 0.00292419 | 394436\|19225\|394432\|54486 |

| ShinyGO Biological Processes Lower expression in HET | | | | | |
|---|---|---|---|---|---|
| Enrichment FDR | nGenes | Pathway Genes | Fold Enrichment | Pathway | Genes |
| 0.0001 | 5 | 139 | 31.49 | Secondary alcohol metabolic process | Soat2 Hsd17b7 Lss Tm7sf2 Scarb1 |
| 0.0001 | 5 | 127 | 34.46 | Cholesterol metabolic process | Soat2 Hsd17b7 Lss Tm7sf2 Scarb1 |
| 0.0001 | 5 | 135 | 32.42 | Sterol metabolic process | Soat2 Tm7sf2 Hsd17b7 Lss Scarb1 |
| 0.0018 | 3 | 36 | 72.95 | Plasma lipoprotein particle | Soat2 Lipg Scarb1 |
| 0.0019 | 3 | 40 | 65.66 | Protein-lipid complex subunit | Soat2 Lipg Scarb1 |
| 0.0021 | 3 | 48 | 54.71 | Secondary alcohol biosynthetic process | Hsd17b7 Lss Tm7sf2 |
| 0.0021 | 3 | 48 | 54.71 | Cholesterol biosynthetic process | Hsd17b7 Lss Tm7sf2 |
| 0.0021 | 5 | 305 | 14.35 | Steroid metabolic process | Soat2 Tm7sf2 Hsd17b7 Lss Scarb1 |
| 0.0024 | 4 | 160 | 21.89 | Steroid biosynthetic process | Tm7sf2 Hsd17b7 Lss Scarb1 |
| 0.0024 | 3 | 54 | 48.63 | Sterol biosynthetic process | Tm7sf2 Hsd17b7 Lss |
| 0.0027 | 5 | 346 | 12.65 | Alcohol metabolic process | Soat2 Hsd17b7 Lss Tm7sf2 Scarb1 |
| 0.0029 | 2 | 9 | 194.53 | High-density lipoprotein particle | Scarb1 Lipg |
| 0.0029 | 3 | 63 | 41.69 | Regulation of plasma lipoprotein | Scarb1 Soat2 Lipg |
| 0.0043 | 8 | 1346 | 5.20 | Lipid metabolic process | Soat2 Tm7sf2 Hsd17b7 Lss Lipg Plch2 Scarb1 Plbd1 |
| 0.0043 | 2 | 12 | 145.90 | High-density lipoprotein particle | Lipg Scarb1 |
| 0.0048 | 3 | 82 | 32.03 | Cholesterol homeostasis | Soat2 Lipg Scarb1 |
| 0.0048 | 2 | 14 | 125.06 | Reverse cholesterol transport | Scarb1 Lipg |
| 0.0048 | 3 | 83 | 31.64 | Sterol homeostasis | Soat2 Lipg Scarb1 |
| 0.0073 | 3 | 97 | 27.07 | Cholesterol transport | Soat2 Scarb1 Lipg |
| 0.0098 | 3 | 109 | 24.09 | Sterol transport | Soat2 Scarb1 Lipg |
| 0.0102 | 5 | 526 | 8.32 | Organic hydroxy compound metabolic | Soat2 Tm7sf2 Hsd17b7 Lss Scarb1 |
| 0.0104 | 2 | 23 | 76.12 | Protein-lipid complex remodeling | Lipg Scarb1 |
| 0.0104 | 2 | 23 | 76.12 | Plasma lipoprotein particle remodeling | Lipg Scarb1 |
| 0.0109 | 2 | 24 | 72.95 | Androgen metabolic process | Hsd17b7 Scarb1 |
| 0.0113 | 2 | 25 | 70.03 | Protein-containing complex | Lipg Scarb1 |
| 0.0150 | 4 | 335 | 10.45 | Lipid catabolic process | Lipg Scarb1 Plch2 Plbd1 |
| 0.0150 | 3 | 138 | 19.03 | Alcohol biosynthetic process | Hsd17b7 Lss Tm7sf2 |
| 0.0150 | 5 | 609 | 7.19 | Lipid biosynthetic process | Tm7sf2 Hsd17b7 Lss Lipg Scarb1 |
| 0.0175 | 2 | 34 | 51.49 | Steroid hormone biosynthetic process | Hsd17b7 Scarb1 |
| 0.0175 | 3 | 152 | 17.28 | Lipid homeostasis | Soat2 Lipg Scarb1 |
| **MGI** | | | | | |
| Enrichment FDR | nGenes | Pathway Genes | Fold Enrichment | Pathway | Genes |
| 6.00E-05 | 4 | 50 | 70.03 | MP:0005278 abnormal cholesterol | Soat2 Lipg Scarb1 Saa2 |
| 6.00E-05 | 3 | 13 | 202.02 | MP:0003980 increased circulating | Lipg Scarb1 Saa2 |
| 0.00134 | 2 | 5 | 350.16 | MP:0000184 abnormal circulating HDL | Lipg Scarb1 |
| 0.00948 | 4 | 226 | 15.49 | MP:0001552 increased circulating | Soat2 Lipg Scarb1 Snap25 |
| 0.01082 | 2 | 17 | 102.99 | MP:0002818 abnormal dentin | Ssu2 Runx2 |
| 0.01536 | 4 | 285 | 12.29 | MP:0005178 increased circulating | Lipg Scarb1 Saa2 Snap25 |
| 0.02292 | 3 | 156 | 16.83 | MP:0001556 increased circulating HDL | Soat2 Lipg Scarb1 |
| 0.02292 | 2 | 35 | 50.02 | MP:0002919 enhanced paired-pulse | Snap25 Syn1 |
| 0.02292 | 1 | 1 | 875.40 | MP:0002897 blotchy skin | Snap25 |
| 0.02292 | 1 | 1 | 875.40 | MP:0030521 abnormal cervical loop | Runx2 |

| ShinyGO Biological Processes Higher in HET | | | | | |
|---|---|---|---|---|---|
| Enrichment FDR | nGenes | Pathway Genes | Fold Enrichment | Pathway | Genes |
| 0.0065 | 13 | 1043 | 4.13 | Response to organic cyclic compound | Pgr Cdh1 Arg1 Ca2 Postn Padi2 Aqp8 Ptgs2 Lum Gjb2 Ugt1a1 Cd36 Ryr3 |
| 0.0065 | 11 | 722 | 5.05 | Cytokine production | Cxcl17 Cd36 Rnf128 Spon2 Naip5 Postn Ptgs2 Arg1 Lum Cd24 Sftpd |
| 0.0065 | 11 | 715 | 5.10 | Regulation of cytokine production | Cxcl17 Cd36 Rnf128 Spon2 Naip5 Postn Ptgs2 Arg1 Lum Cd24 Sftpd |
| 0.0065 | 14 | 1113 | 4.17 | Tube development | Agr2 Tacstd2 Arg1 Prom1 Ptgs2 Gjb2 Cxcl17 Crlf1 Mecom Slc23a1 Pgr Cd36 Sftpd Cd24 |
| 0.0069 | 2 | 3 | 221.06 | Positive regulation of nephron tubule epithelial cell differentiation | Prom1 Cd24 |
| 0.0069 | 10 | 653 | 5.08 | Anion transport | Cd36 Slc34a2 Atp10b Slc23a1 Arg1 Clca1 Nmur2 Ca2 Emb Aqp8 |
| 0.0069 | 30 | 4823 | 2.06 | System development | Emb Dclk1 Agr2 Cd24 Tacstd2 Cdh1 Arg1 Cdkl1 Cyp26a1 Ca2 Postn Prom1 Ptgs2 Lum Nmur2 Gjb2 Sprr1a Fut9 Cxcl17 Ugt1a1 Crlf1 Hoxa10 Mecom Tshr Alcam Slc23a1 Pgr Cd36 Sftpd Pcp4 |
| 0.0069 | 26 | 3884 | 2.22 | Regulation of biological quality | Cd36 Cp Sftpd Ca2 Slc34a2 F13a1 Slc6a2 Atp10b Ryr3 Klk1b21 Ptgs2 Fam3d Cyp26a1 Postn Prom1 Rnf128 Car12 Nmur2 Ugt1a1 Agr2 Tshr Mecom Cd24 Slc28a3 Cxcl17 Ugt1a6 |
| 0.0069 | 2 | 3 | 221.06 | Glomerular parietal epithelial cell differentiation | Prom1 Cd24 |
| 0.0086 | 14 | 1362 | 3.41 | Cell adhesion | Emb Postn Cdh16 Spon2 Cd24 Tacstd2 Cdh1 Cd36 Arg1 Agr2 Alcam Dpt Fut9 Sftpd |
| 0.0086 | 10 | 745 | 4.45 | Response to bacterium | Naip5 Cd36 Spon2 Cd24 Arg1 Ptgs2 Gjb2 Ugt1a1 Mecom Sftpd |
| 0.0086 | 15 | 1607 | 3.10 | Response to endogenous stimulus | Tshr Pgr Cd36 Cdh1 Arg1 Ca2 Postn Padi2 Slc34a2 Aqp8 Ptgs2 Gjb2 Ugt1a1 Cd24 Ryr3 |
| 0.0086 | 14 | 1374 | 3.38 | Biological adhesion | Emb Postn Cdh16 Spon2 Cd24 Tacstd2 Cdh1 Cd36 Arg1 Agr2 Alcam Dpt Fut9 Sftpd |
| 0.0086 | 2 | 4 | 165.80 | Induction of bacterial agglutination | Spon2 Sftpd |
| 0.0086 | 4 | 72 | 18.42 | Maternal process involved in female pregnancy | Arg1 Ptgs2 Gjb2 Pgr |
| 0.0086 | 22 | 3068 | 2.38 | Cellular response to chemical stimulus | Crlf1 Tshr Pgr Cxcl17 Ugt1a1 Ugt1a6 Cd36 Postn Spon2 Cdh1 Arg1 Agr2 Cyp26a1 Clca1 Padi2 Aqp8 Ptgs2 Gjb2 Pcolce2 Cd24 Ca2 Ryr3 |
| 0.0086 | 19 | 2423 | 2.60 | Cellular response to organic substance | Crlf1 Tshr Pgr Spon2 Cdh1 Cd36 Arg1 Agr2 Postn Padi2 Aqp8 Ptgs2 Gjb2 Ugt1a1 Pcolce2 Cd24 Ca2 Cxcl17 Ryr3 |
| 0.0086 | 9 | 595 | 5.02 | Cellular response to organic cyclic compound | Pgr Cdh1 Arg1 Padi2 Aqp8 Ptgs2 Gjb2 Ugt1a1 Ryr3 |
| 0.0086 | 7 | 337 | 6.89 | Renal system development | Tacstd2 Cyp26a1 Ca2 Prom1 Crlf1 Mecom Cd24 |
| 0.0090 | 2 | 5 | 132.64 | Protein citrullination | Padi1 Padi2 |
| **Jensen Tissues** | | | | | |
| Enrichment FDR | nGenes | Pathway Genes | Fold Enrichmen | Pathway | Genes |
| 1.7127E-05 | 8 | 174 | 15.25 | Stromal cell | Hoxa10 Cdh1 Postn Pgr Alcam Prom1 Mmp3 Ptgs2 |
| 3.6332E-05 | 4 | 18 | 73.69 | Glandular epithelium | Ptgs2 Hoxa10 Cdh1 Pgr |
| 1.4112E-04 | 13 | 852 | 5.06 | Adult | Cp Ptgs2 Hpgds Mmp3 Gjb2 Slc6a2 Alcam Prom1 Cdh1 Pgr Ugt1a1 Hoxa10 Postn |
| 9.4370E-04 | 3 | 15 | 66.32 | Uterine epithelium | Hoxa10 Cdh1 Pgr |
| 1.8695E-03 | 5 | 118 | 14.05 | Root nodule | Hpgds Cdh1 Pgr Galnt3 Tshr |
| 3.0092E-03 | 4 | 70 | 18.95 | Cancer stem cell | Dclk1 Cdh1 Alcam Prom1 |
| 3.0092E-03 | 3 | 26 | 38.26 | Conceptus | Pgr Ptgs2 Hoxa10 |
| 3.9562E-03 | 6 | 251 | 7.93 | Plasma cell | Lum Alcam Cp Pigr F13a1 Postn |
| 3.9562E-03 | 11 | 934 | 3.91 | Immune system | Cp Arg1 Pigr Ptgs2 Hpgds Dpt Mmp3 Alcam Prom1 Emb Tshr |
| 1.2897E-02 | 2 | 10 | 66.32 | Temporomandibular joint | Galnt3 Mmp3 |
| **ChEA Database** | | | | | |
| Enrichment FDR | nGenes | Pathway Genes | Fold Enrichmen | Pathway | Genes |
| 1.86E-05 | 16 | 1050 | 5.053 | SUZ12 18692474 ChIP-Seq MEFs Mouse | Crlf1 Ptgs2 Tacstd2 Cd24 Slc34a2 Tmprss2 Ca2 Galnt3 Cyp26a1 Gjb2 Slc6a2 Alcam Prom1 Pgr Hoxa10 Tshr |
| 2.17E-05 | 17 | 1271 | 4.435 | ESR1 22446102 ChIP-Seq UTERUS Mouse | Alcam Prom1 Pgr Postn Nmur2 Lum Padi2 Padi1 Arg1 Pcolce2 Dclk1 Tmprss2 Slc6a2 Emb Slc28a3 Ryr3 Car12 |
| 7.45E-05 | 18 | 1600 | 3.730 | SUZ12 18692474 ChIP-Seq MESCs Mouse | Crlf1 Cd24 Ca2 Cyp26a1 Gjb2 Alcam Prom1 Pgr Hoxa10 Ppp2r2c Ptgs2 Tacstd2 Slc34a2 Tmprss2 Galnt3 Slc6a2 Car12 Tshr |
| 7.65E-05 | 18 | 1635 | 3.651 | SUZ12 18974828 ChIP-Seq MESCs Mouse | Crlf1 Cdkl1 Cd24 Ca2 Cyp26a1 Gjb2 Alcam Prom1 Pgr Ugt1a1 Hoxa10 Ppp2r2c Tacstd2 Pcolce2 Slc34a2 Tmprss2 Galnt3 Slc6a2 |
| 0.0091 | 11 | 1000 | 3.648 | SUZ12 16625203 ChIP-ChIP MESCs Mouse | Tacstd2 Cd24 Slc34a2 Tmprss2 Galnt3 Cyp26a1 Gjb2 Slc6a2 Pgr Hoxa10 Ppp2r2c |
| 0.0091 | 19 | 2612 | 2.412 | MTF2 20144788 ChIP-Seq MESCs Mouse | Crlf1 Cd24 Cyp26a1 Gjb2 Alcam Prom1 Pgr Ugt1a1 Hoxa10 Cdh16 Ppp2r2c Tacstd2 Dclk1 Slc34a2 Tmprss2 Galnt3 Slc6a2 Rnf128 Tshr |
| 0.0158 | 11 | 1103 | 3.307 | RNF2 18974828 ChIP-Seq MESCs Mouse | Crlf1 Tacstd2 Cd24 Ca2 Galnt3 Cyp26a1 Prom1 Pgr Ugt1a1 Hoxa10 Ppp2r2c |
| 0.0158 | 11 | 1103 | 3.307 | EZH2 18974828 ChIP-Seq MESCs Mouse | Crlf1 Tacstd2 Cd24 Ca2 Galnt3 Cyp26a1 Prom1 Pgr Ugt1a1 Hoxa10 Ppp2r2c |
| 0.0187 | 10 | 959 | 3.458 | TP53 20018659 ChIP-ChIP R1E Mouse | Crlf1 Cd24 Spon2 Ggta1 Cyp26a1 Gjb2 Alcam Klhdc7a Hoxa10 Ppp2r2c |
| 0.0224 | 10 | 995 | 3.333 | JARID2 20064375 ChIP-Seq MESCs Mouse | Crlf1 Tacstd2 Cd24 Galnt3 Cyp26a1 Gjb2 Slc6a2 Pgr Hoxa10 Ppp2r2c |
